# Augmentation of a neuroprotective myeloid state by hematopoietic cell transplantation

**DOI:** 10.1101/2023.03.10.532123

**Authors:** Marius Marc-Daniel Mader, Alan Napole, Danwei Wu, Yohei Shibuya, Alexa Scavetti, Aulden Foltz, Micaiah Atkins, Oliver Hahn, Yongjin Yoo, Ron Danziger, Christina Tan, Tony Wyss-Coray, Lawrence Steinman, Marius Wernig

## Abstract

Multiple sclerosis (MS) is an autoimmune disease associated with inflammatory demyelination in the central nervous system (CNS). Autologous hematopoietic cell transplantation (HCT) is under investigation as a promising therapy for treatment-refractory MS. Here we identify a reactive myeloid state in chronic experimental autoimmune encephalitis (EAE) mice and MS patients that is surprisingly associated with neuroprotection and immune suppression. HCT in EAE mice leads to an enhancement of this myeloid state, as well as clinical improvement, reduction of demyelinated lesions, suppression of cytotoxic T cells, and amelioration of reactive astrogliosis reflected in reduced expression of EAE-associated gene signatures in oligodendrocytes and astrocytes. Further enhancement of myeloid cell incorporation into the CNS following a modified HCT protocol results in an even more consistent therapeutic effect corroborated by additional amplification of HCT-induced transcriptional changes, underlining myeloid-derived beneficial effects in the chronic phase of EAE. Replacement or manipulation of CNS myeloid cells thus represents an intriguing therapeutic direction for inflammatory demyelinating disease.

## Introduction

Multiple sclerosis (MS) is the most common inflammatory neurological disorder in young adults, leading to demyelination and progressive disability.^1^ Despite a number of available disease modifying therapies (DMTs), many patients will continue to accumulate neurological disabilities, with a particular risk for patients with early and highly active MS.^2, 3^ In addition, current treatments aim to reduce relapses in MS whereas there is a lack of therapy for progressive disease in which compartmentalized inflammation and ongoing neurodegeneration present challenging obstacles for treatment.^4^ Hence, there is an urgent need for more efficacious and durable therapies, which could be possibly met by cell-based therapies.^5^ Indeed, there is growing evidence for the effectiveness of autologous hematopoietic cell transplantation (AHCT) in MS.^6–10^ Multiple open label studies and trials in selected patient cohorts showed remarkable clinical benefits, which are currently being evaluated in larger clinical trials.^2^

Traditionally, AHCT is preceded by a conditioning regimen, which varied in its intensity between published studies from nonmyeloablative or intermediate-intensity regimens^6, 7, 9, 10^ to higher-intensity regimens including busulfan.^8^ Following AHCT, there are substantial long-term immunological changes in the peripheral blood, such as the development of a renewed T cell repertoire with broader clonal diversity.^11–14^ While these findings explain systemic immunomodulatory mechanisms underlying AHCT- associated remission, especially regarding the lymphocyte compartment, little is known about therapy-induced changes within the central nervous system (CNS), which potentially involves both resident and infiltrating myeloid cell populations. Notably, prior studies have demonstrated that busulfan-conditioned hematopoietic cell transplantation leads to CNS engraftment of donor derived myeloid cells in murine models.^15, 16^ To what extent AHCT might lead to replacement of CNS-resident myeloid cells in neuroinflammatory disease and if this mechanism conveys any of the therapeutic effects remains unknown.

Based on insights gained from animal models of neuroinflammation and demyelination, most notably from experimental autoimmune encephalomyelitis (EAE), which has served as an experimental model for MS for decades and enabled the preclinical discovery of numerous MS treatment strategies,^17^ it is acknowledged that cells of the CNS including the astrocyte, oligodendrocyte, and myeloid lineage have relevant roles in the pathophysiology of neuroinflammation, exhibit broad reactive changes, and can be even potential contributors to a disease-promoting or reparative environment.^18–21^ Particularly the role of microglia is complex, displaying the ability to promote both inflammation and remyelination.^22–24^ Single cell RNA sequencing technology allowed an in-depth depiction of the vast transcriptional alterations in acute EAE, however, the late chronic stages of the disease are less well illuminated.^25, 26^

In this study, we sought to investigate the CNS-specific mechanistic basis of AHCT-induced remission with the ultimate goal to assess the cellular effects of syngeneic bone marrow transplantation (BMT) in general and CNS-resident myeloid cells in particular in the chronic phase of EAE.

## Results

### NucSeq reveals an expansion and prominent transcriptional activation of spinal cord myeloid cells in chronic EAE

In order to assess the unbiased transcriptional responses in CNS cell types in chronic EAE with and without BMT we conducted a comprehensive Single Nuclear Sequencing (NucSeq) experiment on the thoracolumbar segment of a total of 14 spinal cords representing (i) a healthy control group (*Control*), (ii) an EAE-only group (*EAE*), (iii) EAE animals with busulfan-conditioned BMT using donor cells from Ubc-GFP mice (*EAE-BMT*), and (iv) EAE animals with BMT followed by transient pharmacological CSF1R inhibition with PLX5622 to increase microglia replacement with grafted myeloid cells (*EAE-BMT+PLX*).^16^ For other experiments we also included a control group of EAE mice that were only treated with PLX5622 for the same 10-day time period (*EAE-PLX*) (**Fig. 1a**). We used a direct, MOG peptide immunization-based EAE model which reliably induced chronic demyelinating lesions with characteristic immune cell infiltrations primarily consisting of Iba1-positive myeloid cells (**Fig. 1b**). In order to characterize the tissue in the chronic phase of EAE we collected the samples 3 months after EAE induction.

**Figure 1:**
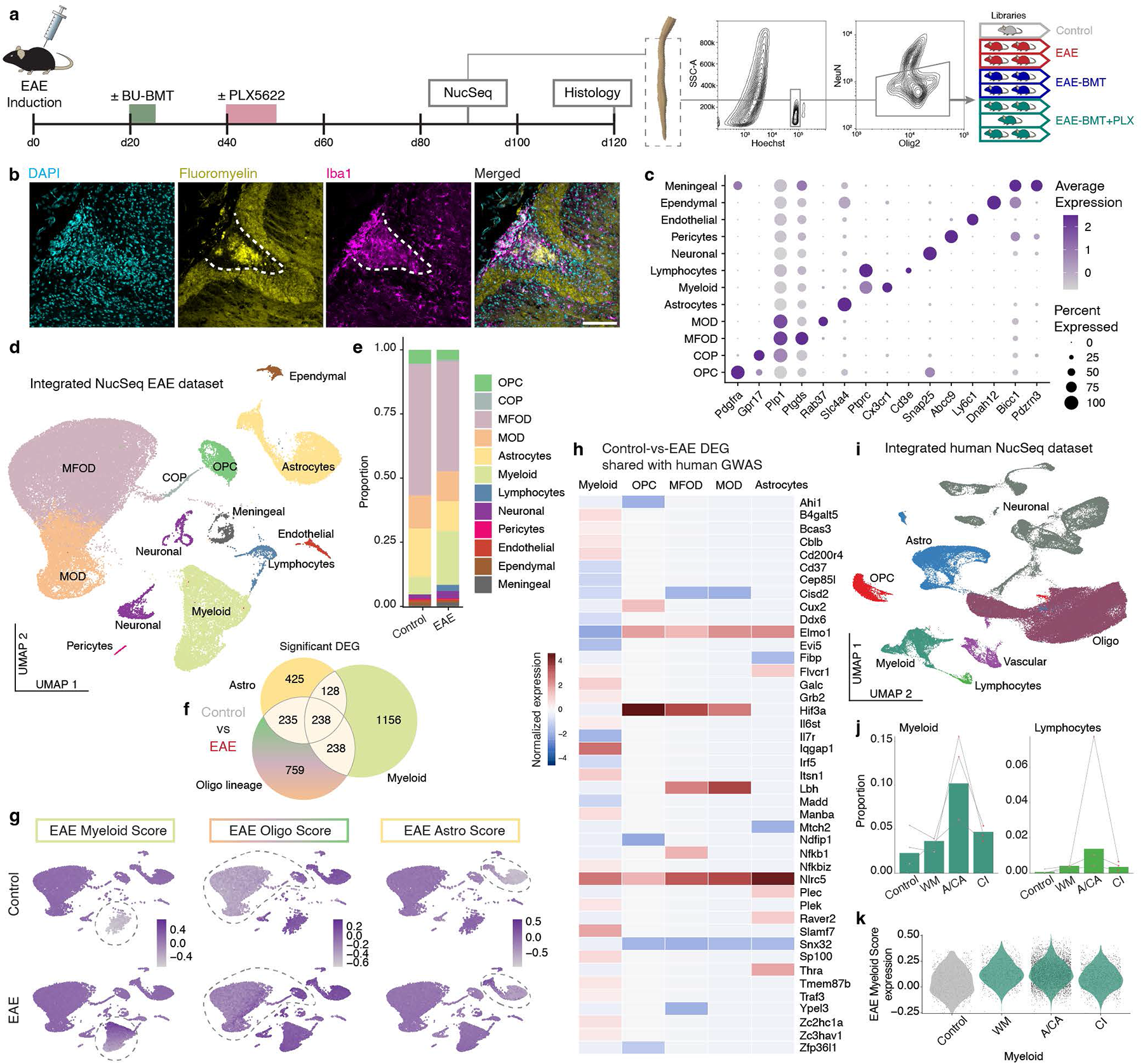
Myeloid expansion and transcriptional activation are features of chronic EAE and MS. (**a**) Experimental design and timeline. Representative image of the lumbothoracic section of the spinal cord used for NucSeq. Contour plots show representative gating for flow-cytometric isolation of NeuN- negative nuclei. Pictogram shows the number of pooled animals per NucSeq library. BU-BMT, busulfan conditioned bone marrow transplantation. EAE, experimental autoimmune encephalomyelitis. NucSeq, single nucleus RNA sequencing. (**b**) Representative image of a lesion in the dorsal column of an *EAE* animal featuring immune infiltration and demyelination. Scale bar 100µm. (**c**) Annotation and canonical marker genes of main clusters. COP, committed oligodendrocyte precursor. MFOD, myelin forming oligodendrocytes. MOD, mature oligodendrocytes. OPC, oligodendrocyte precursor cells. (**d**) Uniform manifold approximation and projection (UMAP) of 68,593 nuclei integrated from all 8 libraries with annotation of main cell clusters. (**e**) Distribution of cell populations between *Control* and *EAE* animals. (**f**) Venn diagram of differentially expressed genes (DEG) of myeloid, astrocyte (Astro), and oligodendrocyte (Oligo) lineage between *Control* and *EAE* condition with an adjusted p value (padj) of <0.05. (**g**) Disease scores were calculated via the VISION^27^ pipeline using gene signatures based on common or cell type-specific DEGs. Score expression changes are shown between *Control* and *EAE* groups. (**h**) Forty-three significant DEGs shared with genes of a multiple sclerosis genome wide association study (GWAS)^29^ are demonstrated with their expression profile over different cell populations. Expression values (average log2 fold-change) are normalized by columns. If no significant DEG was present for a cell type, the value 0 was applied. (**i**) UMAP of 132,425 nuclei integrated from three human NucSeq datasets with annotation of main cell cluster groups. (**j**) Relative proportion of the myeloid and lymphocyte cell clusters. Dots represent different original datasets. A/CA, active / chronic active lesion; CI, inactive lesion; WM, white matter. (**k**) Expression of the EAE Myeloid Score in the myeloid cell population of the human NucSeq dataset. Expression between Control and MS groups differs significantly (p < 2.2e-16, Mann–Whitney U test).

To achieve higher representation of nuclei of immune and oligodendrocyte lineage cells (OLC) we enriched nuclei with low expression of the neuronal marker NeuN by fluorescence activated nuclei sorting. Nuclei derived from animals with similar clinical scores within one experimental group were pooled and a total of 8 sequencing libraries were generated using the 10x platform. After quality control, the integrated dataset consisted of 68,593 nuclei. Yield and quality metrics were comparable between the libraries (**Extended Data Fig. 1a, b**). Based on dimensionality reduction and unsupervised clustering we identified 12 distinct populations of main CNS cell types (**Fig. 1c, d**) with several subclusters (**Extended Data Fig. 1c, d**), which were present in all experimental groups and individual libraries (**Extended Data Fig. 1e**).

First, we sought to characterize the chronic EAE model by comparing *Control* and *EAE* groups. As expected, we observed a substantial change of cellular composition in EAE with a prominent increase of myeloid cells, appearance of lymphocytes and a relative decrease of astrocytes and OLCs (**Fig. 1e**). In addition, the chronic EAE also induced stark transcriptomic changes within cell types of the spinal cord. Differential gene expression analysis between the *Control* and *EAE* groups for each of the 12 cell clusters returned a total of 3369 unique significantly differentially expressed genes (DEGs; adjusted p value (padj) < 0.05). Myeloid lineage nuclei showed the highest number of DEGs (1,760), followed by OLCs (1,470), and astrocytes (1,026) (**Fig. 1f, Extended Data Fig. 1f**). Moreover, about 2/3 of DEGs in myeloid nuclei were unique to myeloid cells, about 1/2 of DEGs in OLCs were OLC-restricted, and only about 2/5 of astrocyte DEGs were astrocyte-specific (**Fig. 1f**). These results illustrate a prominent role of myeloid cells in this chronic phase of EAE.

In line with these data, as further detailed below, we confirmed an overall increase of Iba1-positive myeloid cells in *EAE* by histological analysis (**Fig. 2a, f**). Remarkably, this was not merely based on myeloid infiltrates in demyelinated lesions but also observed in seemingly non-affected CNS tissue depicting a higher overall density of ramified cells.

**Figure 2:**
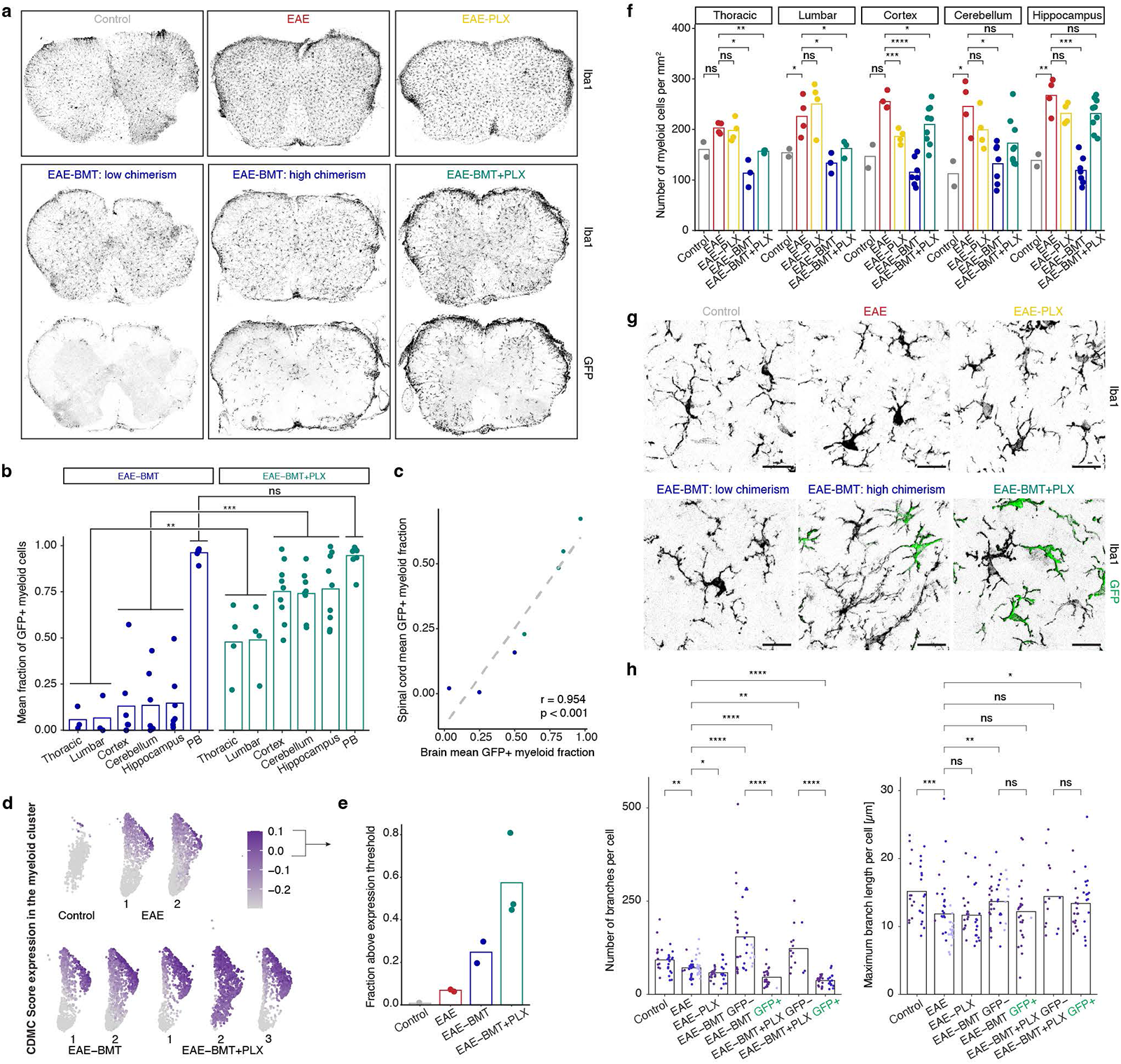
BMT changes the density and morphology of CNS myeloid cells in chronic EAE. (**a**) Representative images of lumbar spinal cord sections demonstrating Iba1+ myeloid cell distribution as well as engraftment of donor derived GFP+ cells. Brightness/contrast was adjusted individually per image for illustrative reasons. (**b**) Chimerism of donor derived GFP+ cells within the myeloid cluster (CNS tissue based on Iba1+ immunofluorescence; peripheral blood (PB) based on CD45+CD11b+ flow cytometry) for different tissues and compared between the *EAE-BMT* and *EAE-BMT+PLX* conditions. Replicates for brain and PB: n=7 and n=9 for *EAE-BMT* and *EAE-BMT+PLX*, respectively. Mean values based on a total number of 136 brain regions of interest (ROIs). Spinal cords do not include tissue used for NucSeq, number of replicates: n=3 and n=4 for *EAE-BMT* and *EAE-BMT+PLX*, respectively, with mean values based on a total of 35 sections. Mann–Whitney U test; ns: p>0.05, *: p≤0.05, **: p≤0.01, ***: p≤0.001. (**c**) Pearson correlation coefficient (r) and scatter plot demonstrate the correlation between brain and spinal cord GFP+ chimerism. Regression line based on linear model. (**d**) A gene signature for CDMCs was based on previously published 1296 DEGs between CDMCs and endogenous microglia^16^ and a score was calculated with the VISION^27^ pipeline. Expression in the myeloid cluster is demonstrated. (**e**) The fractions of nuclei in the myeloid cluster with a CDMC score expression of > - 0.05 are shown. Bars represent the group mean. (**f**) Quantification of ramified Iba1+ cell density. Dots represent the average of multiple measurements per animal. Bars represent the mean of all animals per condition. A total number of 78 spinal cord ROIs and 221 brain ROIs were assessed. t-test; ns: p>0.05, *: p≤0.05, **: p≤0.01, ***: p≤0.001, ****: p≤0.0001. (**g**) Representative immunofluorescent images of ramified myeloid cells in the spinal cord. Brightness/contrast was adjusted individually per image for morphological assessment due to differing Iba1 intensity between conditions. Scale bar is 20µm. (**h**) Morphological analysis of ramified myeloid cells of the spinal cord in different conditions. Bars represent the group mean. Each dot represents one cell, dot colors represent different animals. Mann–Whitney U test; ns: p>0.05, *: p≤0.05, **: p≤0.01, ***: p≤0.001, ****: p≤0.0001.

Myeloid cells demonstrated a prominent enrichment for pathways associated with lipid metabolism featuring the upregulation of genes like *Apoe*, *Abca1*, or *Soat1* (**Extended Data Fig. 2a, b**). Other upregulated pathways included transmembrane receptor protein tyrosine kinase signaling, cell-cell signaling, and defense response. Astrocytes showed a downregulation of energy-related pathways associated with oxidative phosphorylation in combination with an upregulation of inflammatory pathways (**Extended Data Fig. 2a, c**). The latter was driven by genes including *Camk1d*, *C4b*, *A2m*, or *Nlrc5*. OLCs demonstrated a downregulation of cytoplasmatic translation (particularly in myelin forming oligodendrocytes (MFODs)) and biosynthetic pathways (particularly in oligodendrocyte precursor cells (OPCs) and mature oligodendrocytes (MOD), **Extended Data Fig. 3a, b**). OPCs also presented with a disrupted expression of genes associated with OPC differentiation, proliferation, and migration like *Pdgfra*, *Sox8*, *Sox10*, *Fgfr2*, or *Vegfa.* This downregulation was accompanied by an upregulation of inflammatory pathways similar to the astrocytic response, driven by genes including *C4b*, *Apoe*, *Apod*, or *Stat3*. We histologically confirmed the increased expression of complement component 4 (C4) in chronic EAE (**Extended Data Fig. 3c**). Focusing on the main cell populations of the dataset, we used DEG derived gene signatures specific for the myeloid, oligodendrocyte, and astrocyte lineages as well as overlapping genes (**Fig. 1f**) to calculate EAE scores, which showed either cell type-specific or common expression changes between *Control* and *EAE* (**Fig. 1g**).^27, 28^

Here we provide a transcriptional characterization of the chronic stage of EAE, adding to the limited molecular knowledge of this translationally relevant phase of the disease model. To facilitate easy access and usability for the research community we have generated a web application featuring our NucSeq dataset organized by cell type and condition (https://werniglab.shinyapps.io/scthi/).

### CNS myeloid expansion and transcriptional alterations are shared features of chronic EAE and MS

To explore the translational relevance of our EAE gene expression data, we compared DEGs of the main cell populations with large scale MS genome-wide association studies (GWAS) data.^29^ Of 203 human MS susceptibility genes, we matched 197 mouse homologues of which 43 (21.8%) were significantly differentially expressed in either the myeloid, oligodendrocyte, and/or astrocyte lineage (**Fig. 1h**). Myeloid cells were again the cell type with most matching DEGs, with *Iqgap1* representing the gene with the strongest myeloid specific expression change. In addition, within OLC, *Hif3a* demonstrated the highest upregulation, particularly prominent for OPCs, while *Nlrc5* was the strongest overexpressed gene in astrocytes.

To further investigate similarities of the chronic EAE model with human MS pathobiology, we integrated three independent human NucSeq datasets comprised of post-mortem brain tissue from progressive MS and control patients, which have previously been reported.^30–32^ The integrated dataset consisted of 132,425 nuclei. While we observed differences in the total number of nuclei and features of the parent datasets (**Extended Data Fig. 4a, b**), clustering of the integrated dataset delineated major CNS populations which were mostly represented by all three parent datasets with, however, varying frequencies (**Fig. 1i, Extended Data Fig. 4c-e**). Despite this heterogeneity in overall cell type representation, we observed an increase in myeloid and lymphoid cells in all three parent datasets, which was most prominent in active/chronic active (A/CA) lesions (**Fig. 1j**) similar to chronic EAE (compare to **Fig. 1e**). Remarkably, 583 DEGs (55.3%) of the human dataset were shared with DEGs in chronic EAE (2886 human orthologs identified out of 3369 DEGs between murine control and EAE). Similar to our observations in mice, myeloid cells were again among the cells with highest number of significant DEGs between disease and control conditions in the human datasets (**Extended Data Fig. 4f**). Notably, myeloid cells showed an increase in the human ortholog of the EAE Myeloid Score, which was prominent for white matter and A/CA but less in chronic inactive (CI) lesions (**Fig. 1k**).

Collectively, we identify cell-type dependent expression changes of genes in chronic EAE with relevance in MS as previously demonstrated by GWAS studies, and furthermore confirm an expansion and transcriptional activation of myeloid cells in progressive MS, particularly in chronic active lesions, thus suggesting translational relevance of the chronic EAE model regarding myeloid pathobiology.

### BMT in EAE normalizes myeloid cell density and induces a bi-modal myeloid cell morphology

Given the prominent EAE-associated changes in both gene expression and cellular abundance of immune cells, we next explored if BMT would impact the CNS myeloid cell compartment. Busulfan-conditioned BMT in EAE animals (*EAE-BMT*) lead to robust donor chimerism in the peripheral blood and myeloid infiltrates at demyelinated lesions but only low-level engraftment of transplanted circulation-derived myeloid cells (CDMCs) in both brain and spinal cord (**Fig. 2a, b, Extended Data Fig. 5a**). We and others previously reported that transient CSF1R inhibition after BMT can substantially increase the incorporation of such CDMCs into the brain of healthy mice.^15, 16, 33–35^ In order to interrogate the role of bone marrow-derived myeloid cells incorporating into the CNS in chronic EAE, we therefore sought to accomplish a higher donor-derived chimerism within the CNS. Indeed, a 10- day PLX5622 treatment after BMT (*EAE-BMT+PLX*) lead to a significant increase in engraftment efficiency of CDMCs in the CNS (**Fig. 2a, b, Extended Data Fig. 5a**). Of note, the chimerism in the spinal cord was consistently lower than in the brain but markedly increased in the *EAE-BMT+PLX* group (**Fig. 2b, c**). However, brain GFP+ chimerism of myeloid cells was more variable and overall lower than previously reported in healthy mice.^16^ Our NucSeq data also confirmed increased myeloid cell replacement in the *EAE-BMT+PLX* condition. A CDMC expression score based on previous expression data (**Fig. 2d**)^16^ exhibited a strong increase in *EAE-BMT+PLX* mice (**Fig. 2e**).

Quite unexpectedly, BMT greatly affected the cell-type composition in chronic EAE mice. The EAE-induced increase in myeloid density in spinal cord was normalized in both BMT groups (**Fig. 2f**). In the brain, which is less affected in this model, only the *EAE-BMT* group showed reduced myeloid cell numbers (**Fig. 2f**). Furthermore, we observed a change in myeloid morphology in association with both disease and intervention (**Fig. 2g**). As expected, in chronic EAE Iba1+ myeloid cells were less ramified as compared to healthy control mice, and slightly less ramified as compared to EAE mice treated transiently with PLX5622 (**Fig. 2h**). Engrafted GFP+ CDMCs were even less ramified in line with previous findings.^16, 33, 35^ Unexpectedly, however, the remaining endogenous CNS-resident GFP-negative microglia showed a pronounced increase in complexity in both BMT conditions, exceeding the ramification in healthy mice. Thus, BMT has a profound effect on spinal cord-resident microglia in chronic EAE. Similar morphological changes were observed in myeloid cells of the brain cortex (**Extended Data Fig. 5b, c**).

Together, these findings demonstrate that BMT globally impacts CNS myeloid cells not only by the integration of donor derived myeloid cells but also by altering the distribution and morphology of endogenous microglia.

### BMT and microglia replacement lead to clinical improvement, reverse oligo- and astrocytic transcriptional changes, but further promote the myeloid transcriptional response to chronic EAE

We next aimed to explore the functional implication of BMT (*EAE-BMT*) and high-efficiency microglial replacement (*EAE-BMT+PLX*) in chronic EAE. The EAE model led to initial peak clinical severity ∼15 days post-induction with a similar peak clinical score between all experimental groups before any interventions (**Fig. 3a, b**). Animals of the *EAE* group continued with severe neurological symptoms over the chronic course with almost all animals presenting with complete hind leg paralysis at 3 months. We observed clinical improvement starting in close temporal relationship to BMT on day 25, resulting in a significantly reduced clinical score on day 40. The median clinical score of the *EAE-BMT* group remained reduced in the later phase but with increasing interindividual variability. The high-efficiency microglial replacement group (*EAE-BMT+PLX*) led to a more robust clinical improvement, that is preventing mortality and clinical scores >3, and resulting in a continued significantly reduced overall clinical score in the *EAE-BMT+PLX* group up to 3 months. We did not detect sustained clinical benefits in the experimental group receiving the 10-day PLX5622 diet without BMT, which served as a control for the effect of transient microglia depletion. These results point to a critical contribution of CNS myeloid cells to the clinical benefit following BMT in the chronic phase of EAE.

**Figure 3:**
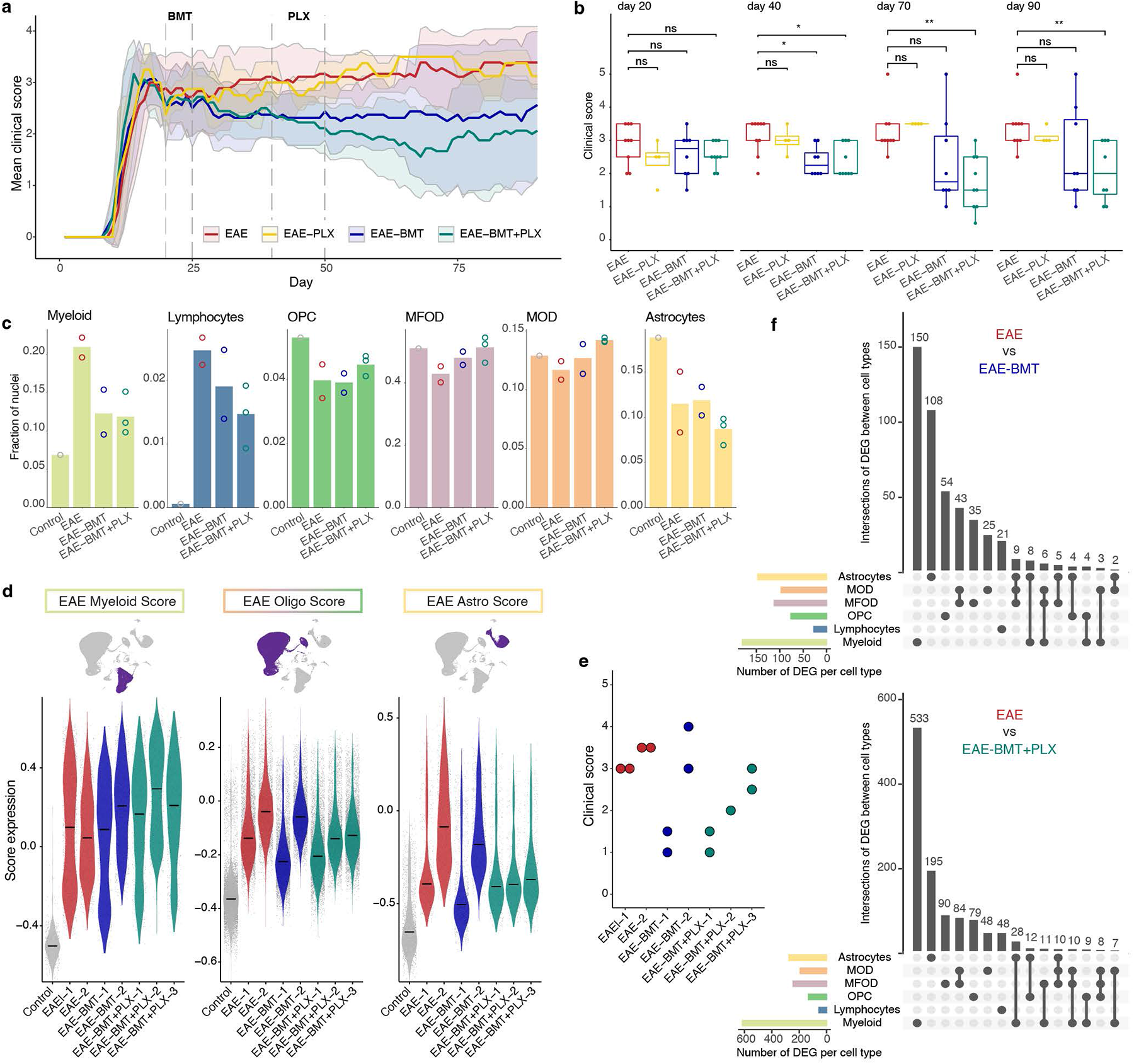
BMT and microglia replacement enhance the myeloid transcriptional response and improve clinical outcome. (**a**) Mean clinical score ± standard deviation over an observation period of 90 days. Thirty animals derived from two independent cohorts were included (*EAE* n=9, *EAE-BMT* n=8, *EAE-BMT+PLX* n=9, *EAE- PLX* n=4). Two animals showed EAE related mortality before completion of the observation period (*EAE* (day 68) and *EAE-BMT* (day 66)). One animal of the *EAE-BMT+PLX* group was sacrificed on day 76 due to non-EAE related reasons (with clinical score of 1.5) and was excluded from the analysis after this timepoint. (**b**) Boxplots show the distribution of clinical scores for the indicated timepoints. Mann– Whitney U test; ns: p>0.05, *: p≤0.05, **: p≤0.01. (**c**) Fractions of nuclei of main cell types in different conditions. Bars represent condition means, dots individual libraries. (**d**) Expression changes of EAE- scores for main cell types between different libraries. Black crossbars represent the median. (**e**) Clinical score of EAE animals used for different NucSeq libraries. (**f**) Upset plots show significant DEGs (padj < 0.05) for main cell types between the *EAE* and *EAE-BMT* or *EAE-BMT+PLX* groups. The top 15 intersections are illustrated.

Next, we assessed cell type representations in EAE and BMT treatment using our NucSeq dataset. Corroborating our previous histological assessment (**Fig. 2f**), we observed an increase in the nuclear fraction of myeloid cells from the *Control* to *EAE* group which was reduced in both *BMT* groups (**Fig. 3c**). Similarly, the EAE-induced increase in lymphocyte nuclei was reduced following BMT. Instead, OPCs, MFOD and MOD were relatively increased in the treatment groups at the expense of astrocytes (**Fig. 3c**) compatible with enhanced myelination explaining the observed clinical improvement (see below and **Fig. 5a-d**).

When comparing the “EAE-scores” for each cell type (established in **Fig. 1g**), we observed an overall reduction of the Oligodendrocyte and Astrocyte scores following BMT again suggesting a partial reversion of EAE-induced transcriptional changes in these two cell types (**Fig. 3d**). Reassuringly, the gene signature scores for these cell types paralleled the degree of paralysis within experimental groups (**Fig. 3e**). Surprisingly, though, the EAE score of myeloid cells behaved in the opposite way: rather than reduced, the score was further increased in both treatment groups (**Fig. 3d** left). In line with the aggregated expression changes, we also detected a relevant number of significant DEGs induced by the interventions (**Extended Data Fig. 6a**). Across all 12 cell clusters we observed 715 total and 542 unique DEGs between the *EAE* and *EAE-BMT* groups, with most DEGs being cell type-specific (**Fig. 3f**). The number of DEGs was further increased between *EAE* and *EAE-BMT+PLX* groups with 1586 total and 1237 unique genes indicating again the prominent role of myeloid cells towards normalizing EAE-induced transcriptional changes in neural cells. Many genes associated with EAE enriched pathways (**Extended Data Fig. 2, 3**) followed the direction of the EAE-scores (**Extended Data Fig. 6b-d**). This applied in particular to an upregulation of lipid metabolism genes in myeloid cells and a reduction in inflammatory pathway genes in astrocytes and OLC.

Collectively, clinical improvement associated with BMT was accompanied with an overall reduced expression of EAE-specific gene signatures in OLC and astrocytes in contrast to an increased expression in the myeloid lineage. These changes were further enhanced after augmented integration of CDMCs in the *EAE-BMT+PLX* group, suggesting a certain protective role of the myeloid response in chronic *EAE*, that can be further augmented by manipulation of the myeloid compartment.

### CNS myeloid cells shift away from a homeostatic state towards transcriptional profiles associated with *injury-responsive* and *disease-associated microglia* in chronic EAE

Given the apparent prominent role of myeloid cells, we next sought to explore deeper the specific changes within this cluster. Subcluster analysis delineated nine myeloid subclusters, in which the expression of canonical microglia-specific genes like *Tmem119*, *Siglech*, *P2ry12*, and *Hexb* was strongest in subclusters M.1 to M.3, while more macrophage- and CDMC-specific genes like *Ms4a7* and *Lyz2* were primarily expressed in M.6 to M.9 (**Fig. 4a, b**).^16, 36, 37^ Despite overall prominent transcriptional changes and the different ontology of microglia and CDMCs, it was surprising to observe that all nine subclusters were present in all conditions albeit in different representations and with intra-cluster transcriptional changes (**Fig. 4c, Extended Data Fig. 7a**). Under the *Control* condition, about 75% of all myeloid cells localized to clusters M.2 and M.5, both of which were reduced in EAE and treatment groups and showed a reduction of homeostatic microglia marker genes such as *P2ry12*, *Siglech*, and *Tmem119* (**Fig. 4c, Extended Data Fig. 7b**). Expression of the CDMC Score was most prominent in clusters M.6 and M.9 (**Fig. 4a**). Immunofluorescence analysis identified CD206, a known marker for border-associated macrophages, as a reliable additional marker for GFP+ CDMCs (**Fig. 4d**). Indeed, *Mrc1* (the gene coding for CD206) demonstrated the highest expression in cluster M.6 (**Fig. 4a**). Accordingly, only few *Mrc1*+ positive cells were detected in nuclei of the *Control* condition, compatible with the low frequency of border associated macrophages in normal mice (**Extended Data Fig. 7c**).^36^

**Figure 4:**
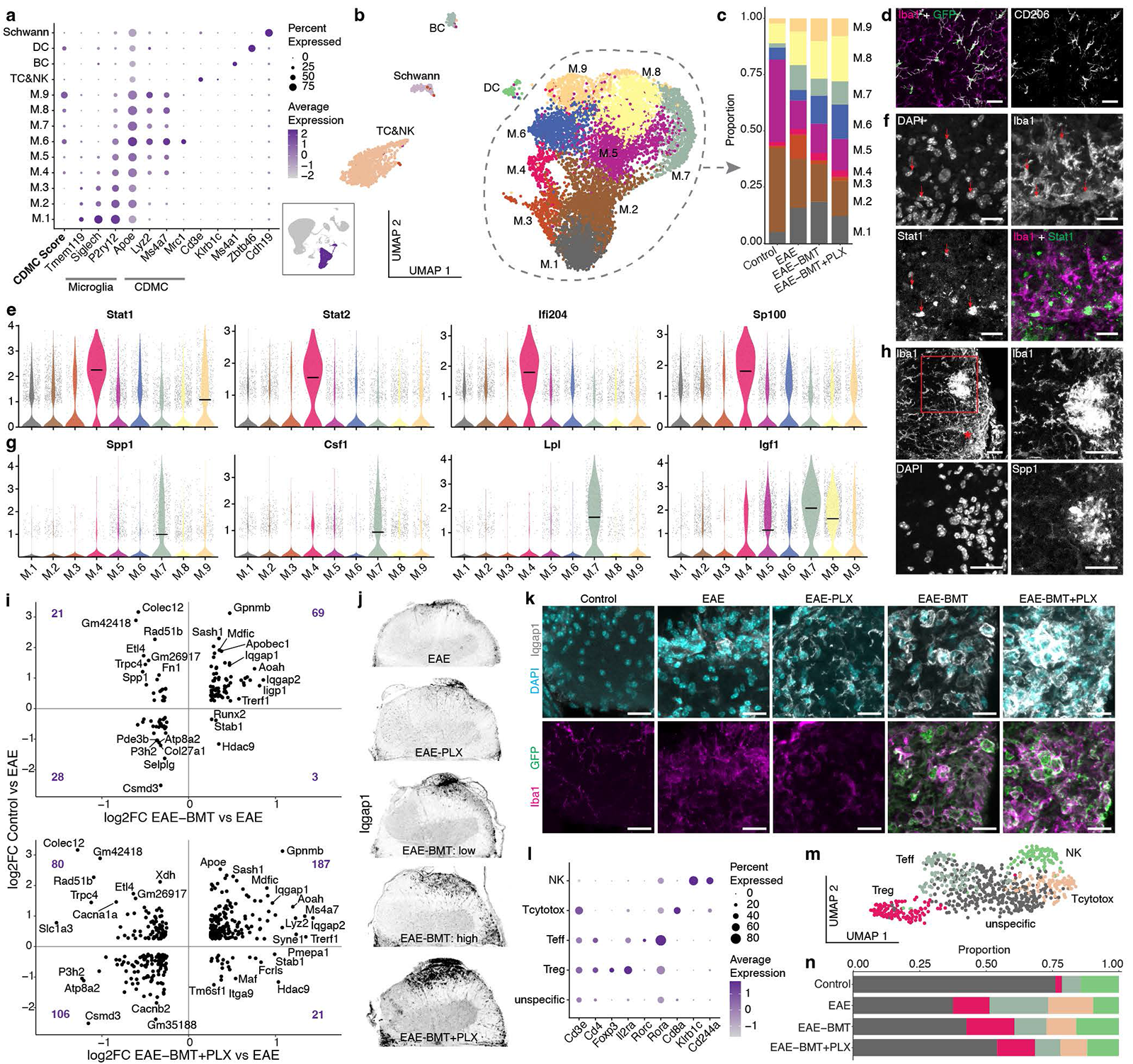
BMT modulates the myeloid and lymphoid response to EAE. (**a**) Subcluster analysis was performed on the immune cell clusters of the integrated main dataset. The dot plot demonstrates the resulting subclusters with an expression profile of canonical marker genes and CDMC Score. BC, B cells. DC, dendritic cells. M, myeloid. NK, natural killer cells. TC, T cells. (**b**) UMAP plot of 10,314 nuclei of the immune cluster. (**c**) Bar chart showing the shifted distribution of myeloid cell subclusters between different conditions. (**d**) Immunofluorescent stain for CD206 (*Mrc1*) in the spinal cord of an *EAE-BMT* animal. Scale bar is 30µm. (**e**) Violin plots of selected genes associated with subcluster M.4 and the interferon pathway. Black crossbars represent the median. (**f**) Immunofluorescent stain for Stat1 in the spinal cord white matter of an *EAE* animal. Red arrows indicate nuclear and paranuclear Stat1 signal colocalizing with Iba1. Scale bar is 20µm. (**g**) Violin plots of selected DAM (disease-associated microglia) genes associated with subcluster M.7. Black crossbars represent the median. (**h**) Representative immunofluorescent image of Osteopontin (Spp1) positive myeloid cells with strong Iba1 signal forming a cluster adjacent to an inflammatory lesion (asterisk) in the spinal cord of an *EAE* animal. Rectangle indicates magnified area with cell cluster. Scale bar is 30µm. (**i**) Scatterplots show the relationship of significant DEGs (padj < 0.05 for both conditions) of disease- associated (*Control vs EAE*) and treatment-associated (*EAE vs EAE-BMT* and *EAE-BMT+PLX*, respectively) conditions of the myeloid cluster. Numbers represent the DEGs per quadrant. Differences between quadrants have been tested with the Pearson’s Chi-squared test with Yates’ continuity correction: p = 2.3e-10 (*EAE-BMT*), p < 2.2e-16 (*EAE-BMT+PLX*). (**j**) Representative immunofluorescent images of hemi spinal cords stained for Iqgap1 including *EAE-BMT* animals with low and high GFP+ chimerism. (**k**) Immunofluorescent stain for Iqgap1 in the spinal cord white matter. Scale bar is 20µm. (l) Marker gene expression of different T cell populations. (**m**) UMAP of the T cell subcluster. (**n**) Distribution of T cell populations between different conditions.

Besides the overall reduction in the expression of homeostatic myeloid genes in EAE, we observed distinct disease-specific gene signatures in myeloid subclusters. Cluster M.4 demonstrated an overexpression of several previously reported injury-responsive microglia (IRM) genes, which were based on a murine injury model of focal demyelination.^38^ IRM genes recognized in M.4, including *Stat1*, *Stat2*, and *Ifi204*, exhibited an association with the interferon pathway, suggesting the continued relevance of interferon signaling in chronic EAE (**Fig. 4e**).^39^ We confirmed the colocalization of *Stat1* in Iba1+ cells in lesions of EAE spinal cords (**Fig. 4f**). Cluster M.7 was almost exclusively present under EAE conditions, thus indicating a disease-specific cell state. Notably, M.7 showed the expression of several genes of disease-associated microglia (DAM), a presumed protective microglia state found in several mouse disease models including for Alzheimer’s disease (**Fig. 4g, Extended Data Fig. 7d**).^40^ Among the overexpressed genes were the critical myeloid cytokine *Csf1*, genes associated with lipid metabolism (*Lpl* and *Cd9*),^41^ and genes also identified in early postnatal axon tract-associated microglia (*Igf1* and *Spp1*).^38^ *Spp1*, which encodes for the secreted glycoprotein Osteopontin, has been implicated in both proinflammatory and neuroprotective microglia functions.^42, 43^ Osteopontin is phosphorylated by the kinase *Fam20c*, which generates the majority of the secreted phosphoproteome and was also found overexpressed in cluster M.7 (**Extended Data Fig. 7d**).^44^ In line with previous reports, *Spp1* expression in M.7 was accompanied with expression of *Itgax*, encoding for the surface protein CD11c (**Extended Data Fig. 7e**).^42^ Immunofluorescence staining revealed Spp1+ cells with high Iba1 expression adjacent to EAE lesions, which were commonly found to form aggregates of multiple cells (**Fig. 4h**). We furthermore observed a shift in the distribution of the remaining clusters, particularly with a rise in the proportion of macrophage-signature clusters M.8 and M.9 in the *EAE* group (**Fig. 4c**). This was further increased in the *EAE-BMT* and *EAE-BMT+PLX* conditions with an additional growth in cluster M.6, while microglia-signature subclusters (M.1 to M.3) decreased. The overall differential gene expression against the *EAE* condition was more pronounced in the *EAE-BMT+PLX* than in the *EAE-BMT* group (**Extended Data Fig. 7f**).

Subclustering of the immune cells of the human NucSeq dataset (**Extended Data Fig. 7g, h**) demonstrated similarly a reduction in a homeostatic myeloid cluster (human M.1) in response to MS (**Extended Data Fig. 7i, j**). For active/chronic active lesions, particularly an increase in the human myeloid cluster M.3 was prominent (**Extended Data Fig. 7i, j**), which was present in all three parent datasets (**Extended Data Fig. 7k**). Importantly, human cluster M.3 showed similarity with the murine EAE-specific cluster M.7, with both demonstrating the expression of DAM genes. Marker genes of human M.3 included the DAM genes *SPP1*, *LPL*, and *APOE* as well as *GPNMB*, and *CD68*, which is associated with phagocytotic activity (**Extended Data Fig. 7l**). A myeloid subpopulation with high expression of *APOE*, *SPP1*, and *LPL* has been independently described in a single cell RNA sequencing study of CD45+ cells isolated from MS patients.^45^

Together this suggests a specific pattern of myeloid activation in the CNS in response to autoimmune neuroinflammation that is partially conserved between chronic EAE and MS.

### A majority of chronic EAE-associated transcriptional changes of the myeloid niche are further enhanced by BMT

Based on the previous observation that the EAE Myeloid Score was further enhanced in both *EAE-BMT* and *EAE-BMT+PLX* conditions (**Fig. 3d**), we sought to further explore this relationship on a DEG level. Indeed, we noticed a positive association between *Control vs EAE* and *EAE vs EAE-BMT* DEGs (**Fig. 4i**). The magnitude of differential expression was further enhanced in the context of *EAE-BMT+PLX*. Importantly, many DEGs were conserved between the two interventions and showed the same direction. We selected *Iqgap1* as a suitable target for validation of these findings given its appearance among human MS-associated GWAS genes (**Fig. 1h**). *Iqgap1* encodes for a cytoplasmic scaffold protein involved in a myriad of cellular functions, including the coordination of signaling events.^46^ Expression in our dataset was increased in association with both EAE and BMT±PLX5622, which was confirmed by immunofluorescent staining (**Fig. 4j, k**). Immuno-reactive areas were primarily detected in white matter areas (**Fig. 4j**). Additionally, we validated a similar expression pattern for *Gpnmb*, which encodes for a glycoprotein associated with an activated microglial population in neurodegenerative disease as well as cancer-associated macrophages (**Extended Data Fig. 8a**).^47, 48^ We observed a staining pattern most likely consistent with previously described endosomal/lysosomal localization.^48^ *Hdac9,* the gene encoding histone deacetylase 9, belonged to a smaller group of DEGs with a negative association between EAE-specific and intervention-specific gene expression changes. Immunostaining demonstrated a nuclear staining pattern of different, also non-myeloid cell populations (**Extended Data Fig. 8b**). In line with our NucSeq data, there was reduced signal intensity of Hdac9 in Iba1+ cells of the *EAE* condition as compared to *Control* samples but an increase from *EAE* to particularly the *EAE- BMT-PLX* condition could be observed.

### BMT is associated with a reduction in cytotoxic and effector T cells in the CNS

We further explored changes within the T cell compartment, given a recent report that microglia directly influence the CNS T cell composition in a relapsing-remitting EAE model, particularly affecting the balance of TH17 effector T cells (Teff) and regulatory T cells (Treg).^49^ Besides an overall decrease in lymphocytes (**Fig. 3c**), subcluster analysis revealed also an effect on T cell composition in association with BMT and further enhancement with microglia replacement (**Fig. 4l-n**). We observed an increase in the Treg/Teff ratio, mainly driven by a reduction in *Rora*+ *Rorc*+ Teff. Moreover, the relative amount of Cd8+ cytotoxic T cells was reduced after BMT. This suggests that BMT drives the CNS T cell compartment away from a proinflammatory towards a more immunosuppressive state.

### BMT induces increased myelination in chronic EAE and reverses EAE-associated transcriptional changes in OLC

Demyelination remained a common histopathological feature in chronic EAE (**Fig. 5a**). The area of demyelination correlated with clinical severity, which was most significant for the cervical and lumbar segments (**Fig. 5b**). Importantly, BMT led to a significant reduction in demyelinated areas in the cervical and lumbar spinal cord, which was further enhanced in the *EAE-BMT+PLX* group (**Fig. 5c**). This histological finding was confirmed by transcriptional analysis as we discovered an upregulation of several key myelination (*Plp1*, *Opalin*, *Mal*, *Mag, Cnp*) and OLC differentiation (*Aspa, Olig1, Slc8a3*) genes among DEGs between the *EAE* and particularly the *EAE-BMT+PLX* groups (**Fig. 5d**). *Slc8a3* (NCX3) has already previously been associated with a protective role in EAE.^50^ *Daam2*, which has been reported to inhibit oligodendrocyte differentiation, was reduced in its expression.^51^ When comparing the DEGs from different conditions, a large proportion of DEGs between EAE and the BMT groups was also among the DEG in chronic EAE compared to normal mice (**Fig. 5e**). Focusing on these DEGs shared between *Control vs EAE* and *EAE vs EAE-BMT/EAE-BMT+PLX*, we observed a striking negative association demonstrating that chronic EAE-induced transcriptional changes are partially reversed by BMT, and further reversed following more efficient microglia replacement in BMT+PLX mice (**Fig. 5f**). This pattern was similar in all OLC differentiation stages but represented by different genes, with e.g., *Hif3a* as a prominent OPC-associated and *Zfand4, Rhoj*, and *Zbtb16* as MOD/MFOD-associated genes. These findings corroborated our earlier observation that overall, the transcriptional EAE oligodendrocyte score decreased following BMT (**Fig. 3d**).

**Figure 5:**
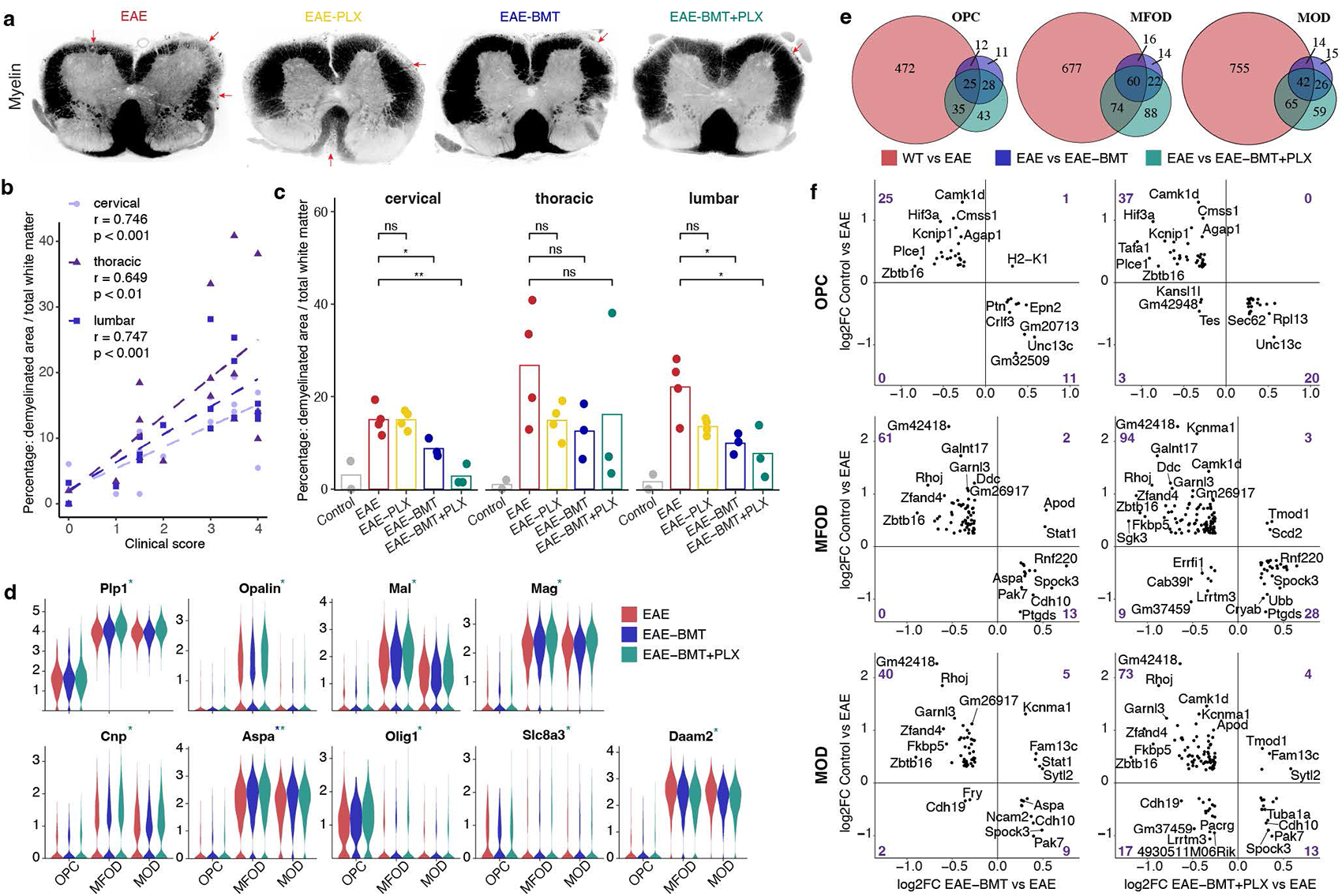
BMT-associated increased myelination is accompanied by partial reversal of the transcriptional response in OLCs. (**a**) Representative fluorescent myelin stain (FluoroMyelin™) of the lumbar spinal cord. Red arrows indicate demyelinated lesions. (**b**) Scatterplot demonstrating the correlation between white matter demyelination and clinical score. Regression lines based on linear models; r = Pearson correlation coefficient. (**c**) Comparison of demyelinated white matter areas between conditions for animals with available spinal cord histology. Dots represent the average of multiple analyzed spinal cord sections, with a median of 4 (range: 2 - 6) sections per anatomical location per animal. Bars represent the mean of all replicates per condition. t-test; ns: p>0.05, *: p≤0.05, **: p≤0.01. (**d**) Violin plots of selected genes associated with myelin formation and oligodendrocyte differentiation/maturation that are differentially expressed between *EAE* and *EAE-BMT* (blue asterisk) or *EAE-BMT+PLX* (green asterisk) conditions (padj < 0.05). (**e**) Venn diagrams of DEGs between selected conditions (padj < 0.05) for main OLC subpopulations. (**f**) Scatterplots show the relationship of significant DEGs (padj < 0.05 for both conditions) of disease-associated (*Control vs EAE*) and treatment-associated (*EAE vs EAE-BMT* and *EAE- BMT+PLX*, respectively) conditions of different OLC subpopulations. Numbers represent the DEGs per quadrant. Differences between quadrants have been tested with the Fisher’s Exact Test: p = 1.4e-08 (*EAE-BMT*) and p = 4.2e-13 (*EAE-BMT+PLX*) for OPC. p = 6.9e-13 (*EAE-BMT*) and p < 2.2e-16 (*EAE- BMT+PLX*) for MFOD. p = 1.2e-05 (*EAE-BMT*) and p = 7.5e-06 (*EAE-BMT+PLX*) for MOD.

### EAE-associated reactive astrogliosis is reduced by BMT and microglia replacement

Reactive astrocytosis has been shown to play an important role in MS pathobiology.^30^ Likewise, we observed an increase in GFAP immunoreactivity in the chronic EAE spinal cord, represented by a denser cell layer in the outer white matter and dispersed ramified cells more distant from the surface (**Fig. 6a**). The elevated GFAP+ area was significantly reduced in both the *EAE-BMT* and *EAE-BMT-PLX* groups, implying amelioration of reactive astrogliosis (**Fig. 6b**). On a single cell level, astrocytes demonstrated two major groups of subclusters with either high expression of genes including *Gpc5* and *Trpm3* (subclusters Astro.1 to Astro.5) or *Glis3*, *A2m*, *Aqp4*, and *Gfap* (subclusters Astro.6 and Astro.7) (**Fig. 6c, d**). Overall, the changes in cellular distribution among these 7 clusters among different conditions were rather subtle (**Fig. 6e**). In line with our histological analysis, clusters Astro.6&7 (characterized by high GFAP expression) increased in EAE and were reduced in the BMT groups. However, all other astrocyte clusters also induced GFAP in chronic EAE and reduced expression again following BMT indicating that all spinal cord astrocyte subtypes become reactive in chronic EAE (**Fig. 6f**). Inspecting additional markers of reactive astrocytes mirrored *Gfap*, such as *Slc1a3, Slc1a2, Stat3, C3, Serpina3n*, and *Vim*, with the latter three being primarily represented in Astro.6+7 (**Fig. 6f**).^52^ *Slc1a3* demonstrated a significantly lower differential expression in *EAE vs EAE-BMT*, while both *Gfap* and *Slc1a3* were significant DEGs for *EAE vs EAE-BMT+PLX*. Similar to expression changes in oligodendroglia (**Fig. 5e**), a majority of DEGs between EAE and BMT groups were also de-regulated in chronic EAE (**Fig. 6g**). Exploring the directionality of those overlapping DEGs, we observed again a negative correlation between EAE-induced changes and BMT-induced changes (**Fig. 6h**). These results illustrate that BMT as well as microglia replacement are able to reverse some of the EAE-induced expression changes, just as we had seen in OLC but not in myeloid cells (**Fig. 4i, 5f**).

**Figure 6:**
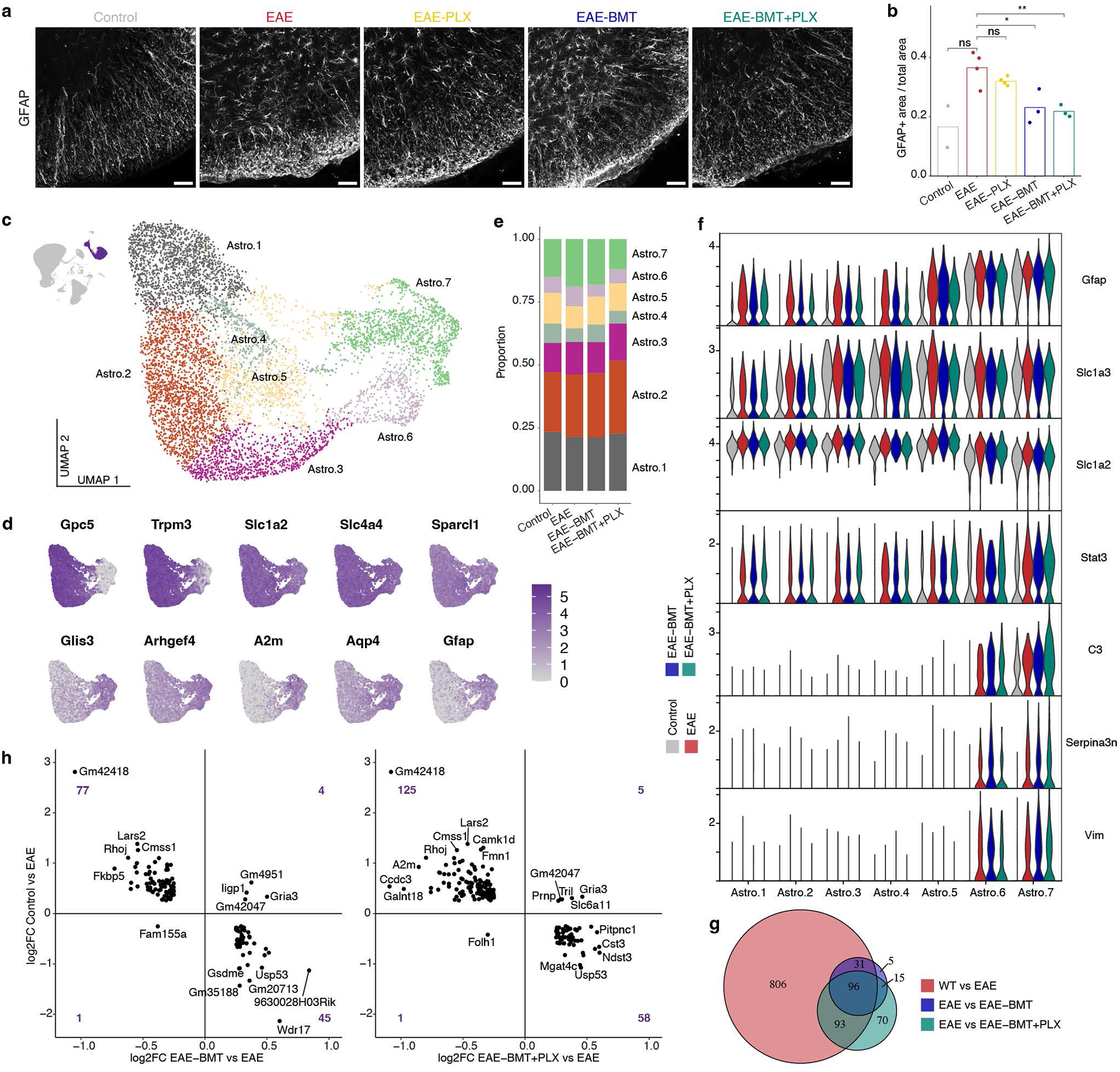
BMT and microglia replacement ameliorate reactive astrogliosis in chronic EAE. (**a**) Immunofluorescent stain for GFAP in the spinal cord. Scale bar is 50µm. (**b**) Quantification of GFAP+ area in the lumbar spinal cord. Dots represent the average of measurements per animal (median number of spinal cord sections per animal = 2). Bars represent the mean of all animals per condition. t-test; ns: p>0.05, *: p≤0.05, **: p≤0.01. (**c**) UMAP plot of the astrocyte cluster consisting of 7745 nuclei. (**d**) Expression of astrocyte subcluster marker genes. (**e**) Bar chart showing the distribution of astrocyte subclusters between different conditions. (**f**) Expression of selected genes associated with reactive astrogliosis. All genes shown are differentially expressed between *Control* and *EAE* (padj < 0.05). Compared to *EAE*, *Slc1a3* and both *Gfap* and *Slc1a3* were differentially expressed versus *EAE- BMT* and *EAE-BMT+PLX*, respectively. (**g**) Venn diagram of DEGs of the astrocyte lineage (padj < 0.05). (**h**) Scatterplots show the relationship of significant DEGs (padj < 0.05 for both conditions) of disease- associated (*Control vs EAE*) and treatment-associated (*EAE vs EAE-BMT* and *EAE-BMT+PLX*, respectively) conditions in the astrocyte cluster. Numbers represent the DEGs per quadrant. Differences between quadrants have been tested with the Pearson’s Chi-squared test with Yates’ continuity correction: p < 2.2e-16 (*EAE-BMT*), p < 2.2e-16 (*EAE-BMT+PLX*).

## Discussion

Here we report a comprehensive dataset addressing the chronic stage of EAE, consisting of 68,593 single-cell level transcriptomes of the murine spinal cord supported by immunohistological and clinical assessment. We found that chronic EAE maintains pathological hallmarks such as demyelinating lesions and chronic immune cell infiltrations which are accompanied by prominent transcriptional changes and subcluster heterogeneity affecting different lineages, of which myeloid cells exhibit the strongest alterations. In order to replicate real-life treatment paradigms and increase translational value, we selected therapy time points after reaching clinical nadir as opposed to prophylactic approaches aimed at preventing EAE induction or early interventions in the acute phase. Consequently, we observed interindividual variability in clinical presentation and gene expression. Despite variability, there were robust changes in EAE-specific DEGs, overlap with MS susceptibility genes (**Fig. 1h**), similarities with the human myeloid response in MS (**Fig. 1j, k, Extended Data Fig. 7l**), as well as detectable effects of experimental interventions. Our characterization of chronic EAE might therefore inform the design and interpretation of future preclinical efforts.

We utilized the chronic EAE model to explore the CNS cell-specific impact of BMT as a model for AHCT in human MS patients. While the therapeutic efficacy of AHCT is currently being validated in clinical trials,^2^ this study provides further evidence of therapeutic efficacy in a preclinical EAE mouse model of a chronic phase of MS. A key finding of this study was that myeloid cells play a prominent and beneficial role in in the chronic phase of the disease, potentially by resolving the consequences of the acute immune attack against myelin.^24^ Following BMT, oligodendroglial and astroglial cells showed a partial reversion of the transcriptional response to EAE – reflecting enhanced remyelination, reduced astrogliosis, and amelioration of paralysis. In stark contrast, the transcriptional response to EAE in spinal cord myeloid cells was further enhanced rather than “normalized” following BMT. To functionally test the role of BMT-derived myeloid cells, we increased their incorporation into the CNS using a recently developed microglia replacement protocol.^15, 16, 33–35^ Increased incorporation of these circulation-derived myeloid cells further improved clinical paralysis and further enhanced the EAE-induced, myeloid transcriptional program. At large, these results strongly suggest that the myeloid response in chronic EAE is at least in part rather neuroprotective, an observation made in the context of remyelination before.^23, 24^

Given that the enhanced microglia replacement protocol involves transient CSF1R inhibition, it can be debated whether the beneficial effect observed in this study is truly based on the cellular properties of CDMCs or rather the combination of two separate effects derived from BMT and PLX5622-mediated CSF1R inhibition. We believe the latter to be unlikely because PLX5622 administration was only transient, of short duration, and is generally followed by rapid repopulation of the myeloid niche.^53, 54^ Moreover, we have included an *EAE-PLX* group to control for PLX5622-specific effects and did not detect clinical improvement. In contrast, long-term inhibition of CSF1R signaling, which is essential for microglial survival, is known to have very different effects as demonstrated in different studies investigating the contribution of CSF1 to EAE pathogenesis.^55–60^ Early and continuous treatment with a CSF1R inhibitor could reduce disease severity in some studies underlining the relevance of myeloid cells in EAE pathobiology.^55–58^ However, drug withdrawal led to a relapse in symptoms suggesting that no sustained modulation of the myeloid niche was achieved thus limiting translational value.^57^ These results, in light of the findings of our study, demonstrate that the role of myeloid cells is complex. They suggest that there might be opposing roles of CNS-resident and circulation-derived myeloid subpopulations and pharmacological CSF1R inhibition results in overall clinical improvement in certain stages of disease. Additionally, the role of CNS-myeloid cells may change over time and may be necessary for the initiation of an immune attack against myelin in the CNS early in EAE but then later switch to a restorative function in the chronic phase of the disease as suggested before.^22–24^

Regardless, our model supports the notion that the combination of CSF1R inhibition with HCT might have translational implications, and our data would warrant further exploration of the specific role and mechanisms of replaced myeloid cells in the context of inflammatory CNS disease. Notably, with Pexidartinib (PLX-3397) a first CSF1R inhibitor has already gained FDA approval for oncological application, underlining overall clinical feasibility and relevance of CSF1R inhibition.^61^ Our findings identify manipulation of the myeloid niche as a therapeutic direction for inflammatory demyelinating disease, warranting the exploration of methods enhancing neuroprotective myeloid cell states in MS.

## Methods

### Animal procedures

Female C57BL/6J mice were used for the EAE model (JAX, Strain 000664). Female C57BL/6-Tg(UBC-GFP)30Scha/J mice of 10-12 weeks were used as donor animals for BMT (JAX, Strain 004353). All animal procedures were approved by the administrative panel on laboratory animal care at Stanford University (APLAC21565). Mice were group-housed (up to 5 mice per cage) on a 12h/12h light/dark cycle with water and standard chow ad libitum, unless otherwise described below.

### Experimental autoimmune encephalitis

EAE was induced using the Hooke Kit™ MOG_35-55_/CFA Emulsion PTX (Hooke Laboratories, Inc., EK-2110) according to the manufacturer’s instructions in female C57BL/6J mice at the age of 11 weeks. On day 0, animals received a percutaneous injection of 100µl on two sites (total of 200µl) of the antigen emulsion of MOG_35-55_ in complete Freund’s adjuvant. This was followed by an intraperitoneal injection of 100µl freshly prepared pertussis toxin with a concentration of 200ng/µL in cold PBS after 2 hours. The preparation and injection of pertussis toxin solution was repeated after 24 hours. Injections were carried out under isoflurane sedation. Animals without clinical presentation (clinical score = 0) were excluded from the experiment before the initiation of any therapeutic measures. In the early stages of disease and later in cages with animals of clinical scores ≥3.0, additional gel food was provided (DietGel® Boost, ClearH_2_O, INC.). Additional supportive care was provided to animals with a clinical score >3.0 in the form of subcutaneous saline injection. Animals with mortality before day 50 were excluded from the analysis. The observation period was either three months for animals used for NucSeq or four months for spinal cord histological analysis.

### Clinical scoring

The clinical evaluation was performed blinded regarding the experimental groups. Clinical scoring was applied according to the manufacturer’s instructions (Hooke Kit™, Hooke Laboratories, Inc., EK-2110). In brief, a scale from 0 to 5 with increments of 0.5 was used. Most relevant clinical presentations are: no obvious motor deficits, a limp tail, weakness of hind legs, complete paralysis of hind legs, additional partial front leg paralysis, and death/euthanasia due to severity of symptoms, which correspond to scores of 0, 1, 2, 3, 4, and 5, respectively. A detailed table is provided by the manufacturer (https://hookelabs.com/protocols/eaeAI_C57BL6.html). Animals requiring euthanasia for animal welfare reasons unrelated to EAE severity were excluded from the analysis after the timepoint of euthanasia.

### Busulfan conditioning and bone marrow transplantation

Busulfan (SIGMA-ALDRICH, B2635) conditioning was started after peak disease severity on day 20 after EAE induction. A working solution of 1mg/ml was prepared fresh every day from a DMSO stock solution of 49mg/ml by dilution with PBS. Busulfan was applied intraperitoneally with a daily dose of 25mg/kg over 5 subsequent days. Control animals received injections with PBS. BMT was performed on day 25. Ubc-GFP donor mice were euthanized with CO_2_. Tibias and femurs were pestled in ice-cold PBS to harvest the bone marrow. The cell solution was then filtered through a 70µm cell strainer and spun down for 5min at 350G. The cell pellet was washed one time in ice-cold PBS, cell number was counted, and cells were again resuspended in ice-cold PBS. Preconditioned recipient mice were retro-orbitally injected with 5 million bone marrow cells in 200µl PBS.

### PLX5622 treatment

PLX5622 was provided by Plexxikon Inc. under a material transfer agreement and incorporated to AIN- 76A standard chow by Research Diets Inc. at 1200 parts per million. Mice were treated with PLX5622 or control diet ad libitum from day 41 until day 50 after EAE induction. Chow was placed both in the food reservoir of the cage and on the cage floor to allow for easy access for paralyzed animals.

### Immunofluorescence staining and microscopy

The following primary antibodies were used for immunofluorescence staining: C4 (abcam, ab11863), CD206 (R&D systems, AF2535), GFAP (Invitrogen, 13-0300), GFP (abcam, ab13970), GPNMB (Bioss, BS- 2684R), HDAC9 (abcam, ab109446), Iba1 (FUJIFILM Wako, 019-19741), Iba1 (abcam, ab5076), IQGAP1 (abcam, ab133490), Olig2 (Millipore Sigma, AB9610), Spp1 (Santa Cruz Biotechnology, sc-21742), Stat1 (Cell Signaling, 14994). Tissue collection for immunofluorescent analysis was performed 4 months after EAE induction or in age-matched healthy control animals. Animals were anesthetized (100 mg/kg of ketamin and 10 mg/ kg of xylazine, i.p.), and transcardiac perfusion with ice-cold PBS followed by 4% PFA was performed before harvesting brain and spinal cord. NucSeq animals were sacrificed 3 months after EAE-induction, perfused with ice-cold PBS, and brains were drop-fixed in 4% PFA. All tissues were fixed in 4% PFA for 18 hours at 4°C. Tissue was then moved into a 30% sucrose in PBS solution. Brains were then embedded with Tissue-Tek OCT compound (Sakura Finetek, 4583), and sections of 40μm were obtained using a cryostat (CM 3050S, Leica). After washing with PBS, sections were incubated in IF buffer (PBS with 5% cosmic calf serum and 0.3% Triton X-100 (Sigma-Aldrich)) for 1 hour. If FluoroMyelin Red Fluorescent Myelin Stain (Invitrogen, F34652) was used, sections were incubated with FluoroMyelin 1:300 in PBS for 30min after the blocking step and subsequently washed in PBS. Primary and secondary antibodies were diluted in IF buffer. Primary antibody staining was performed at 4°C overnight. Sections were subsequently washed three times in PBS. Secondary antibody (ThermoFisher) stain was performed at room temperature for 1 hour. Sections were subsequently washed three times in PBS and counterstained with DAPI for 5min. Sections were washed once with PBS and mounted on glass slides with drop ProLong Gold antifade reagent (ThermoFisher, P36930). Images were obtained using a Zeiss LSM710 confocal microscope, Zeiss AxioImager motorized widefield fluorescence microscope, or Leica DMi8 using consistent imaging parameters per stain.

### Image analysis and quantification

Image postprocessing and analyses were performed using ImageJ (NIH, USA) software. Postprocessing workflow and parameters were kept consistent per stain unless otherwise stated. For chimerism and myeloid density, images were taken using a fluorescence microscope. For the brain, 10x images were taken from the cortex, hippocampus, and cerebellum, sampling between 2-4 sections per region. For the spinal cord, tiled images were taken from axial thoracic and lumbar cords. Quantification of chimerism and myeloid density was performed manually using CellCounter plugin. Percent chimerism was determined by GFP+Iba1+ cells divided by total Iba1+ cells. For the density of myeloid cells, total Iba1+ cells were divided by region of interest area. For Fluoromyelin analysis, tiled fluorescent images were taken from axial cervical, lumbar, and thoracic spinal cord section. All stained spinal cords were imaged and analyzed, resulting in between 2-6 sections per replicate per anatomical region. Quantifications were performed blinded. To quantify the percentage of demyelination, boundaries were manually drawn around areas of demyelination at the edge of the spinal cords which was defined as regions with DAPI+Fluoromyelin- staining and divided by total white matter area which was defined by total spinal cord area minus grey matter area. For GFAP analysis, tiled fluorescent images were taken from axial thoracic and lumbar cord. All stained spinal cord sections were imaged resulting in 2- 4 sections per replication per anatomical region. Images were converted into 8-bit grayscale and total GFAP area was obtained using a set lower signal boundary applied using the standard Threshold plugin. GFAP positive area was normalized by dividing the value by the total spinal cord area. For branch analysis, confocal images of microglia/CDMCs were obtained from spinal cord grey matter and cortex at 40x, Z-stack of 25µm at 1µm Z-interval. Brightness and contrast were optimized to allow for visualization of morphology due to variations in Iba1 staining related to the treatment condition. Images were converted into 8-bit grayscale images and processed using standard plugins (i.e. despeckle, FTT bandpass, thresholding, unsharp mask, despeckle, close, skeletonize). Skeletonized images were analyzed using Analyze Skeleton plugin.

### Peripheral blood analysis and flow cytometry

Peripheral blood was collected from the facial vein and treated with 0.5M EDTA. Erythrocytes were removed by 10 minutes of incubation in 2ml ice-cold RBC lysis buffer.^62^ Cells were collected via centrifugation for 10min at 300g. Cells were washed with PBS and then stained for 15min at 4 °C in Flow buffer (PBS with 5% cosmic calf serum) containing the following antibodies: CD11b-PE (Invitrogen, 12-0112-82), CD45-APC (Biolegend, 103112). Stained cells were washed in PBS and resuspended in Flow buffer with DAPI (Sigma, D9542). Flow cytometry was performed on a FACS AriaII. Data analysis was performed using FlowJo software. The percentage of GFP+ cells within the CD45+CD11b+ live population was determined.

### Isolation of nuclei from frozen spinal cord tissue

Tissue collection for the transcriptomic analysis was performed 3 months after EAE induction or in an age-matched healthy control animal. Animals were anesthetized (100 mg/kg of ketamin and 10 mg/ kg of xylazine, i.p.) and transcardially perfused. Spinal cords were harvested by hydraulic expulsion and snap-frozen by submerging for 60 seconds in liquid nitrogen-cooled isopentane. Frozen spinal cords were stored at -80°C until further processing. Single-nuclei preparation was performed as previously described with the following modifications.^63^ The thoracolumbar section of spinal cords were first mechanically dissociated with an ice-cold razor blade in a petri dish with 1ml EZ Prep lysis buffer (Sigma, NUC-101) on ice. Transfer of tissue suspension into a 2ml glass dounce tissue grinder tube (Sigma, D8938). Petri dish was rinsed off with another 1ml EZ buffer, which was added to the grinder tube. Suspension was homogenized by hand 25 times with pestle A followed by 25 times with pestle B while incorporating a 180-degree twist. Tissue homogenate was transferred to a fresh 15ml tube on ice. The grinder tube was rinsed with 2 ml fresh lysis buffer, which was added to the 15ml tube for a total volume of 4 ml. Samples were incubated on ice for 5 minutes. Nuclei were centrifuged at 500 x g for 5 minutes at 4°C, supernatant was removed, and the pellet resuspended with 4 ml EZ lysis buffer and incubated on ice for 5 minutes. Centrifugation at 500 x g for 5 minutes at 4°C was repeated. After removing the supernatant, the pellet was resuspended with 4 ml chilled PBS and filtered through a 35- um cell strainer into a 5 ml round bottom FACS tube (Corning, 352235). Following centrifugation at 300 x g for 10 minutes at 4°C with break 3, supernatant was gently poured out leaving behind the nuclei pellet. The pellet was resuspended in 100µl FC block buffer (BD Biosciences, 553141, 1:50 in FACS buffer) and 1ul recombinant RNase inhibitor (Takara, 2313B) incubated on ice for 5min. FACS buffer consisted of PBS (Gibco, 10010049) containing 1% BSA (Thermo Fisher, BP9700100)). To 50µl of the cell suspension, we added 150µl FACS buffer with 2µl Olig2-Alexa Fluor488 (Sigma, MABN50A4), 1ul NeuN-AlexaFluor647 (Abcam, ab190565), and 1.5ul RNase inhibitor. After 30min incubation covered from light and shaking on ice, 3ml FACS buffer were added. Following centrifugation at 300 x g for 10 minutes at 4°C with break 3, supernatant was gently poured out. The pellet was resuspended in 700µl FACS buffer with 3.5ul recombinant RNase inhibitor and Hoechst dye (Thermo Fisher, H3570, 1:2000) and incubated 4min on ice covered from light. Single nuclei were sorted on a MA900 Multi-Application Cell Sorter (Sony Biotechnology) gating for NeuN negative nuclei. 100,000 single nuclei per sample were collected into 1.5 ml DNA lo-bind tubes (Eppendorf, 022431021) containing 1 ml buffer mix with 800ul PBS, 200ul UltraPure BSA (Thermo Fisher, AM2618), and 5ul RNase inhibitor. If libraries were representing two animals, samples were pooled by collecting identical numbers (2x 50,000) of nuclei into the same sample tube. Collected nuclei were centrifuged at 400 x g for 5 minutes at 4°C with break 2. Supernatant was removed leaving 40µl suspended nuclei. Nuclei were counted using a hemacytometer (Sigma, Z359629-1EA) and assessed for concentration and quality.

### Droplet-based single nuclei RNA sequencing

Reagents of the Chromium Next GEM Single Cell 3’ GEM & Gel Bead Kit v3.1 (10X Genomics, 1000128) were thawed and prepared according to the manufacturer’s protocol. Nuclei and master mix solution was adjusted to target 10,000 nuclei per sample and loaded on a standard Chromium Controller (10X Genomics, 1000204) according to manufacturer protocols. We applied 11 PCR cycles to generate cDNA. Library construction was conducted using Chromium Next GEM Single Cell 3ʹ Library Kit v3.1 (10X Genomics, 1000128). All reaction and quality control steps were carried out according to the manufacturer’s protocol and with recommended reagents, consumables, and instruments. We chose 11 PCR cycles for library generation. Quality control of cDNA and libraries was conducted using a Bioanalyzer (Agilent) at the Stanford Protein and Nucleic Acid Facility. Illumina sequencing of the resulting libraries was performed by Novogene (https://en.novogene.com/) on an Illumina NovaSeq S4 (Illumina). Base calling, demultiplexing, and generation of FastQ files were conducted by Novogene.

### Mapping, integration, quality control, clustering, annotation, and subclustering of NucSeq data

The Cell Ranger (v.6.1.2) analysis pipelines were utilized to align reads to the mm10 reference genome as well as *GFP*, and count barcodes/UMIs. The nucleotide sequence for *GFP* was retrieved by Sanger sequencing of a PCR product generated from genomic DNA of a C57BL/6-Tg(UBC-GFP)30Scha/J mouse with the following primer pair: forward CTTTAGTTGCAAGGGTTTCTG and reverse ATCAACTTTCCTGTGCGTAGG, which was based on previous mapping.^64^ To account for unspliced nuclear transcripts, reads mapping to pre- mRNA were counted. Separate Seurat objects were created per library, including features expressed by at least a minimum of 3 cells and including cells with expression of at least 50 features.^65^ Outliers with a ratio of mitochondrial relative to endogenous RNAs of >5%, homotypic doublets (>6,000 features) and debris (<400 features) were removed. We integrated all eight libraries, using Seurat’s built-in SCTransform^66^ and integration workflow with 2000 genes set as integration features. The integrated dataset was used as input for cell embedding and clustering. A shared-nearest-neighbors graph was constructed using the first 20 PC dimensions before clustering spots using Seurat’s built-in FindClusters function with a resolution of 0.6 and default parameters. UMAPs were calculated using Seurat’s built-in functions, based on the first 20 PC dimensions. Count data was subsequently normalized and scaled to allow for visualization of expression values and differential gene expression analysis. Seurat’s FindAllMarkers function (parameters: only.pos = TRUE, min.pct = 0.15, logfc.threshold = 0.15, assay = ’SCT’) was run to identify cluster markers. Clusters were annotated based on canonical marker genes derived from previous publications.^26, 38, 67–70^ Nuclei cluster were manually inspected for debris and doublet clusters, which were removed for downstream analysis. Cluster expressing more than one cell type-specific marker signature were defined as doublets. For subclustering, new Seurat objects were generated from clusters of the main integrated dataset with Seurat’s subset function. Each new object was reintegrated using the SCTransform based workflow with subsequent clustering as outlined above. Clustering parameters were 15 PC dimensions and a resolution of 0.6 for immune cells (combined main clusters “Myeloid” and “Lymphocytes”, see **Fig. 1d**) and 10 PC dimensions and a resolution of 0.4 for astrocytes. Downstream processing including annotating was performed as described above. Further subclustering for T cells from the immune cell subcluster was performed with 15 PC dimensions and a resolution of 0.6, followed by removal of a small cluster with Schwann cell signature and a cluster with predominant expression of Ribosomal protein L (Rpl) genes. A pseudobulk expression matrix was generated with the built-in Seurat function AverageExpression (slot = “counts“) for visualization in a Shiny app, which can be accessed via https://werniglab.shinyapps.io/scthi/.

### Differential gene expression analysis and Gene Set Enrichment Analysis (GSEA)

Differential gene expression analysis was performed for each of the main 12 clusters of the main integrated NucSeq dataset comparing the different experimental groups. Calculation was based on the MAST algorithm^71^ with a log-fold-change threshold of 0.25, minimum detection fraction of 0.1 and Bonferroni correction of p values. Differentially expressed genes (DEGs) were considered significant with an adjusted p value (padj) of < 0.05. Gene Set Enrichment Analysis (GSEA) was performed using the fgsea package, genes were ranked based on log2 fold-change expression, and pathways with 15 to 500 genes were considered.^72^

### Gene signature score calculation

The generation of gene signature scores was performed using the VISION (v.3.0) package as detailed in the original study, using default settings without pooling and signed signatures.^27^ VISION z-normalizes signature scores with random gene signatures to account for global sample-level metrics.

### Analysis of human single nuclei RNA sequencing data

Human NucSeq datasets were obtained as expression matrices from the UCSC Cell Browser (https://cells.ucsc.edu)73 and were originally generated by Absinta et al. (3 control and 5 MS patients), Jäkel et al. (5 control and 4 MS patients), and Schirmer et al. (9 control and 12 MS patients).^30–32^ For the dataset of Schirmer et al., nuclei with a maximum number of 10,000 features were used and duplicated gene symbols were kept as distinct genes. No further adaptions were made to the other two datasets before integration. We integrated the datasets using Seurat’s built-in SCTransform^66^ and integration workflow with 2000 genes set as integration features. The integrated dataset was used as input for cell embedding and clustering. A shared-nearest-neighbors graph was constructed using the first 20 PC dimensions before using Seurat’s built-in FindClusters function with a resolution of 0.6 and default parameters. UMAPs were calculated using Seurat’s built-in functions, based on the first 20 PC dimensions. Count data was subsequently normalized and scaled to allow for visualization of expression values and differential gene expression analysis. Seurat’s FindAllMarkers function (parameters: only.pos = TRUE, min.pct = 0.15, logfc.threshold = 0.15, assay = ’SCT’) was run to identify cluster markers. Nuclei cluster were manually inspected for doublet clusters, which were removed for downstream analysis. Differential gene expression analysis was based on the MAST algorithm^71^ with a log-fold-change threshold of 0.25, minimum detection fraction of 0.1 and Bonferroni correction of p values. Differentially expressed genes (DEGs) were considered significant with an adjusted p value (padj) of < 0.05. Expression of the EAE Myeloid Score, based on a signed signature of the human orthologs, was calculated from the UMI-scaled expression matrix using the VISION package without pooling and default settings.^27^ Immune cells were subclustered by sub-setting the “Myeloid” and “Lymphocyte” clusters, followed by reintegration and clustering as described above with clustering parameters of 15 PC dimensions and a resolution of 0.4. Subclusters with relevant expression of non-immune cell marker genes were excluded for downstream analysis.

## Data availability

The datasets generated and analyzed in the current study are available from the corresponding author upon request. Averaged gene expression data can be accessed via a web application: https://werniglab.shinyapps.io/scthi/

## Code availability

Computational and statistical analysis have been performed using freely available software packages. Custom code generated in this study is available from the corresponding author upon reasonable request.

## Acknowledgments

We would like to thank all members of the Wernig laboratory, Gordon Wang, Chris Bennett, Bahareh Ajami, Jacob Blum, and Noga Or-Geva for helpful discussions throughout the project, Amy Lang and Madhuri Vangipuram for administrative support, and the FACS core at the Institute for Stem Cell Biology and Regenerative Medicine, and Stanford Neuroscience Microscopy Service (supported by NIH NS069375) for technical support.

## Author contributions

Study concept and design: MM, MW, LS, RD, CT. Animal work: MM, AN, YS, DW, YY, RD, CT. Single nuclei RNA sequencing: MM, AF, MA, OH, TWC. Immunostaining, microscopy and image analysis: DW, AS, MM, AN. Data analysis and interpretation: MM, MW, OH, AN, DW, LS. Drafting and major editing of original manuscript: MM, MW. All authors reviewed, revised, and approved the final version of the paper.

---

**Extended Data Figure 1.**
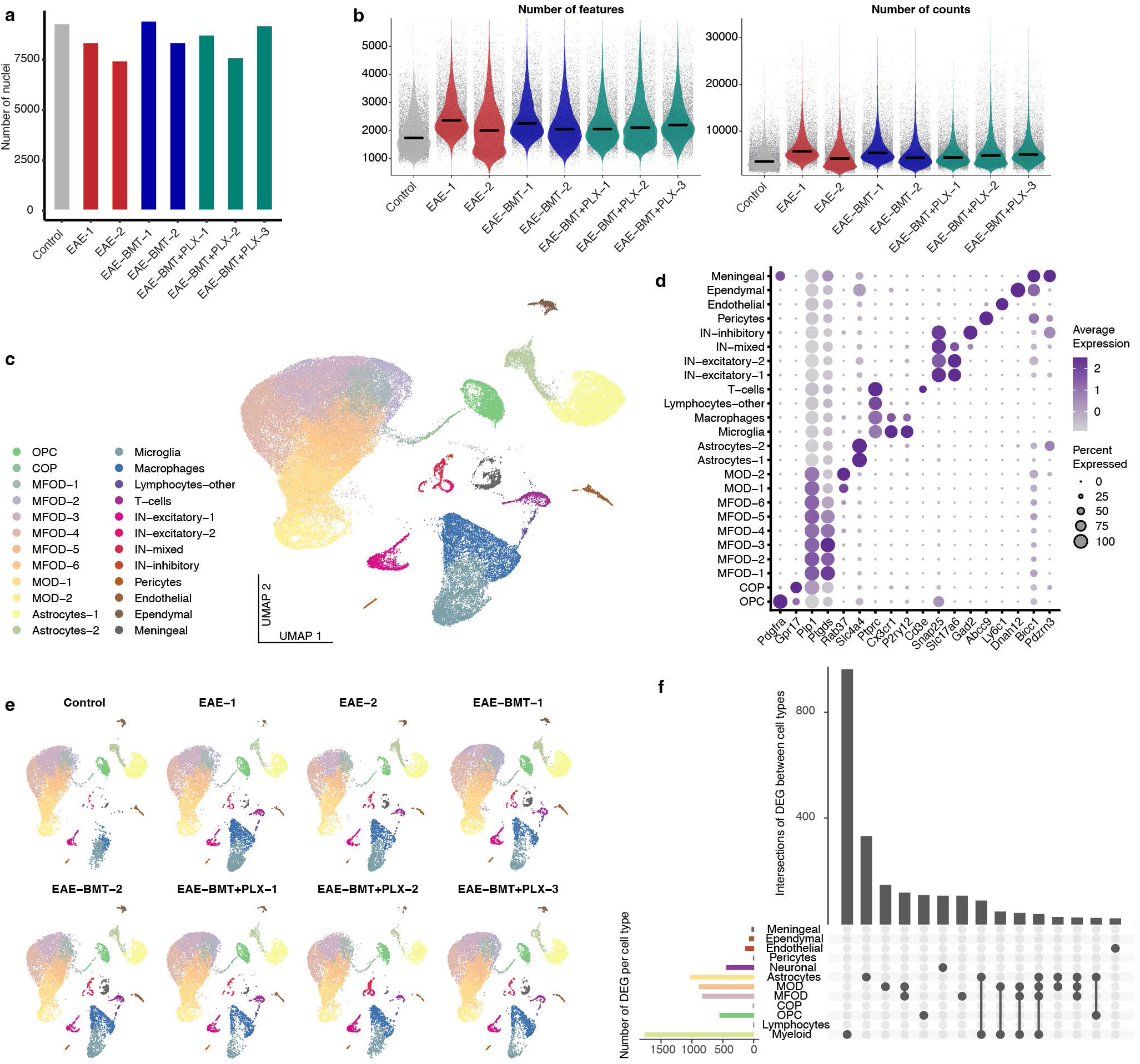
(**a**) Absolute number of nuclei per library. (**b**) Number of features and counts per library. (**c**) Annotated UMAP plot of 68,593 nuclei integrated from all 8 libraries. Clustering based on the first 20 principal components and a resolution of 0.6. (**d**) Dotplot demonstrating annotation and canonical marker genes of clusters. (**e**) UMAPs of individual libraries. (**f**) Upset plot showing significant DEGs (padj < 0.05) for main cell types between the *Control* and *EAE* condition. The top 15 intersections are illustrated.

**Extended Data Figure 2.**
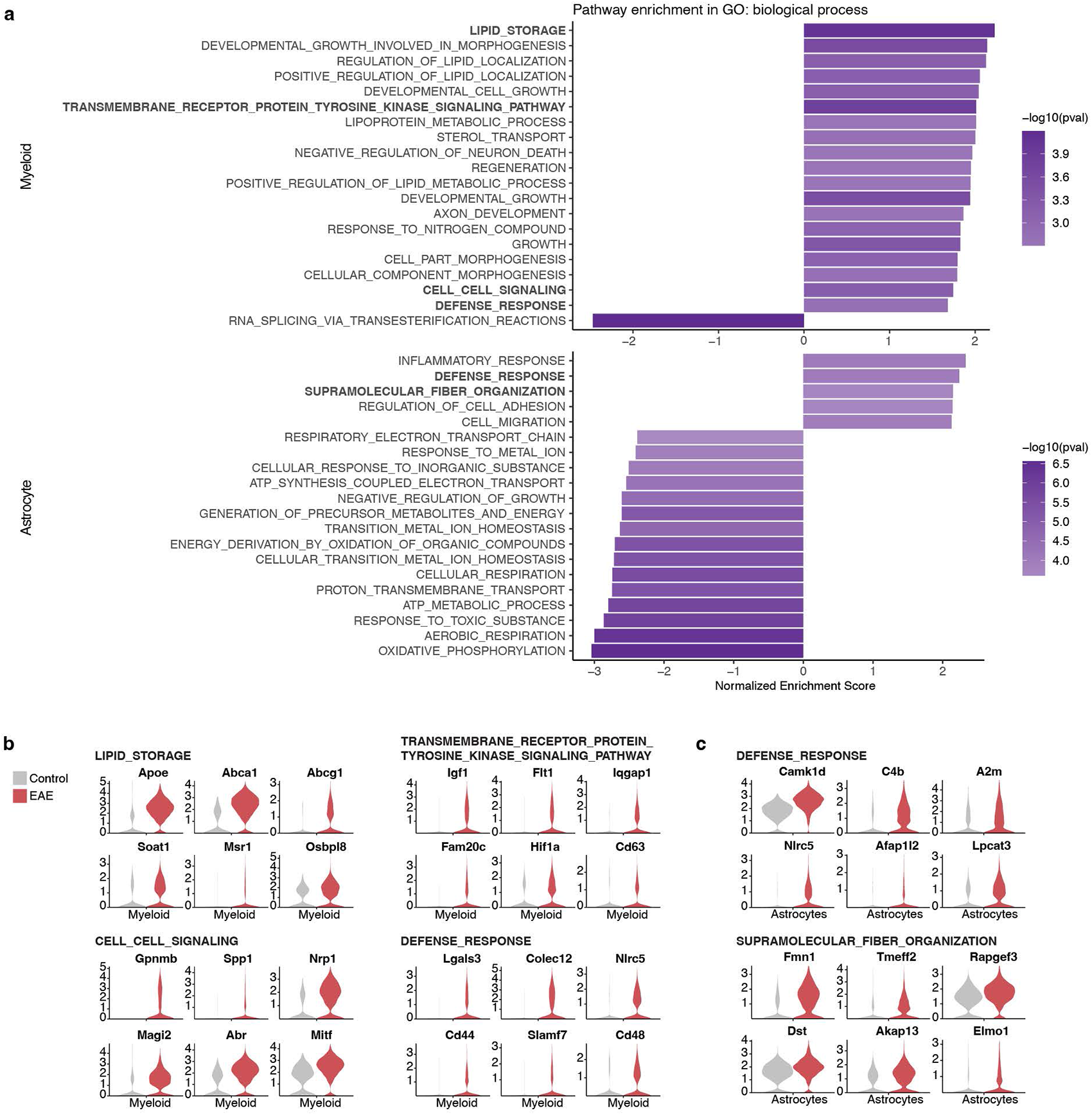
(**a**) GSEA-derived top 20 significantly enriched pathways (*control* vs *EAE*) of the Gene Ontology biological process term for the myeloid and astrocyte clusters. (**b**)+(**c**) Representative genes of selected enriched pathways

**Extended Data Figure 3.**
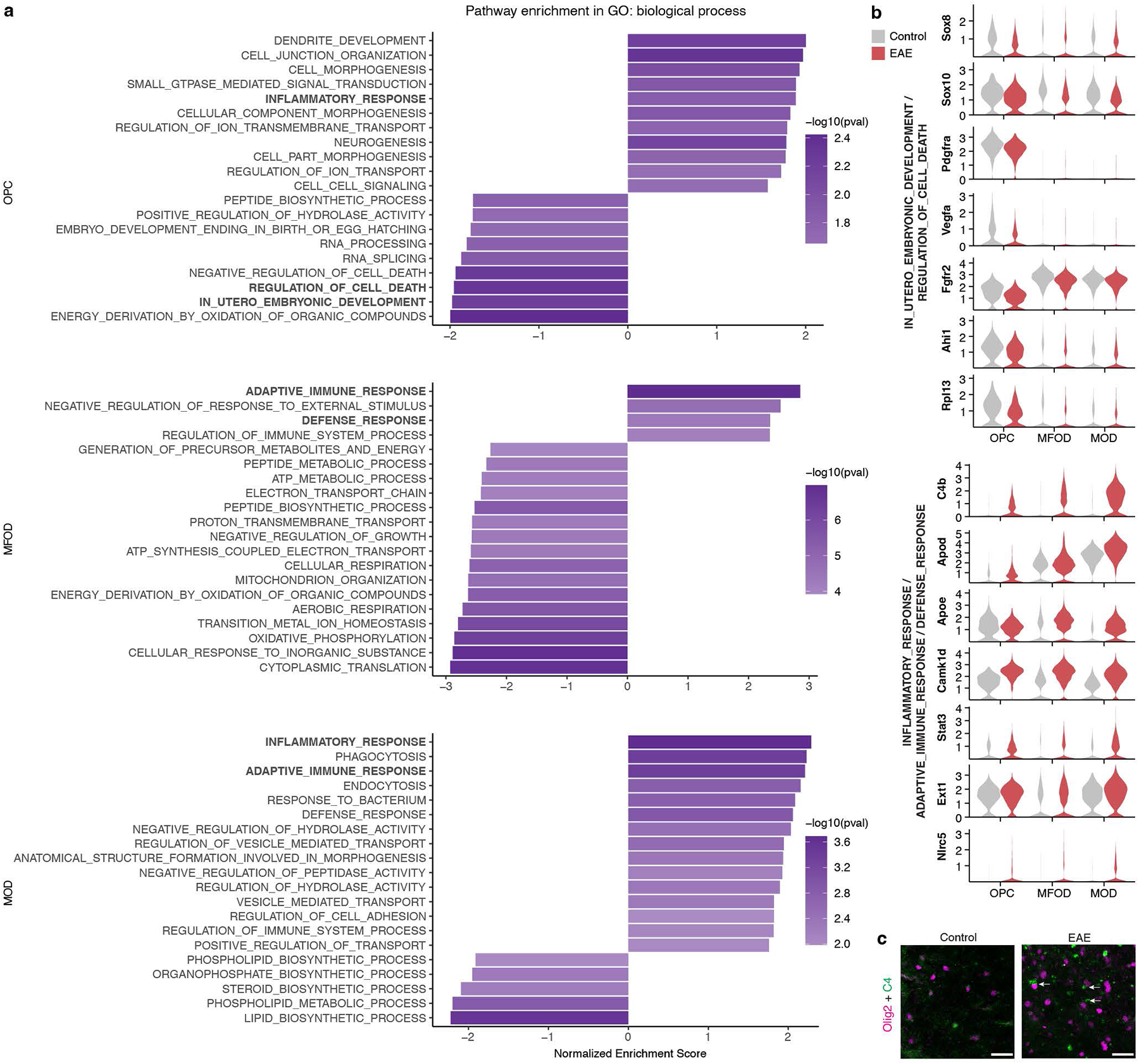
(**a**) GSEA-derived top 20 significantly enriched pathways (*control* vs *EAE*) of the Gene Ontology biological process term for OLC clusters. (**b**) Representative genes of selected enriched pathways. (**c**) Immunofluorescent stain for Olig2 and complement component 4 (C4) in the spinal cord white matter. White arrows demonstrate spatial association between C4 and OLC. Scale bar is 20µm.

**Extended Data Figure 4.**
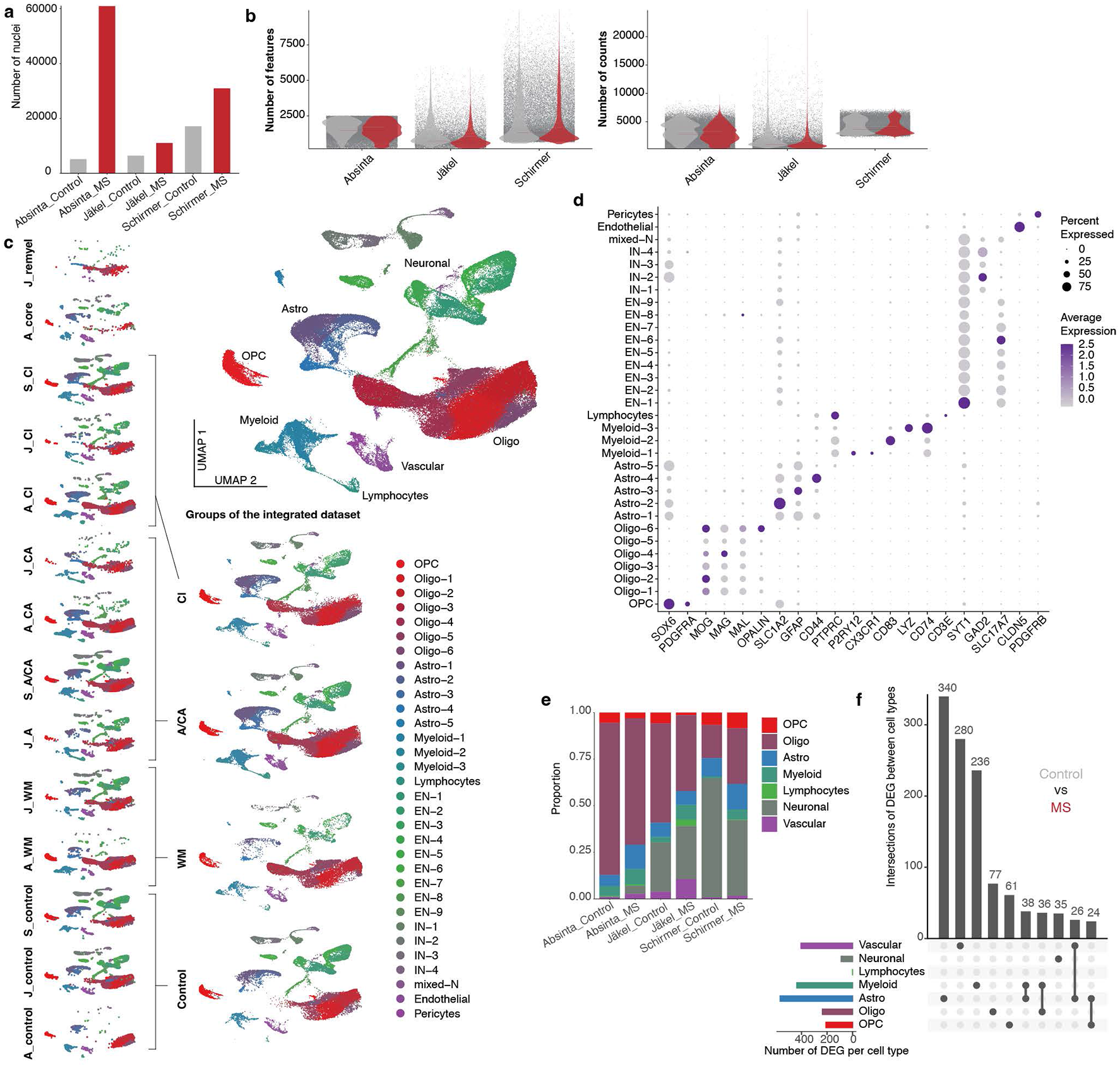
(**a**) Absolute number of nuclei per original dataset. (**b**) Number of features and counts original dataset. (**c**) UMAP plots of 132,425 nuclei integrated from the original datasets. Clustering based on the first 20 principal components and a resolution of 0.6. Lesion-dependent conditions of the original datasets are shown (A_, Absinta; J_, Jäkel; S_, Schirmer; A, active; A/CA, acute/chronic active; CA, chronic active; CI, chronic inactive; remyel, remyelinating; wm, white matter), which were combined into 4 groups: control, active/chronic active (A/CA), chronic inactive (CI), and MS white matter (WM). (**d**) Dotplot demonstrating annotation and canonical marker genes of clusters. (**e**) Relative distribution of main cluster groups between different original datasets and control/MS condition. (**f**) Upset plot showing significant DEGs (padj < 0.05) for main cell types between the *Control* and *MS* condition. The top 10 intersections are illustrated.

**Extended Data Figure 5.**
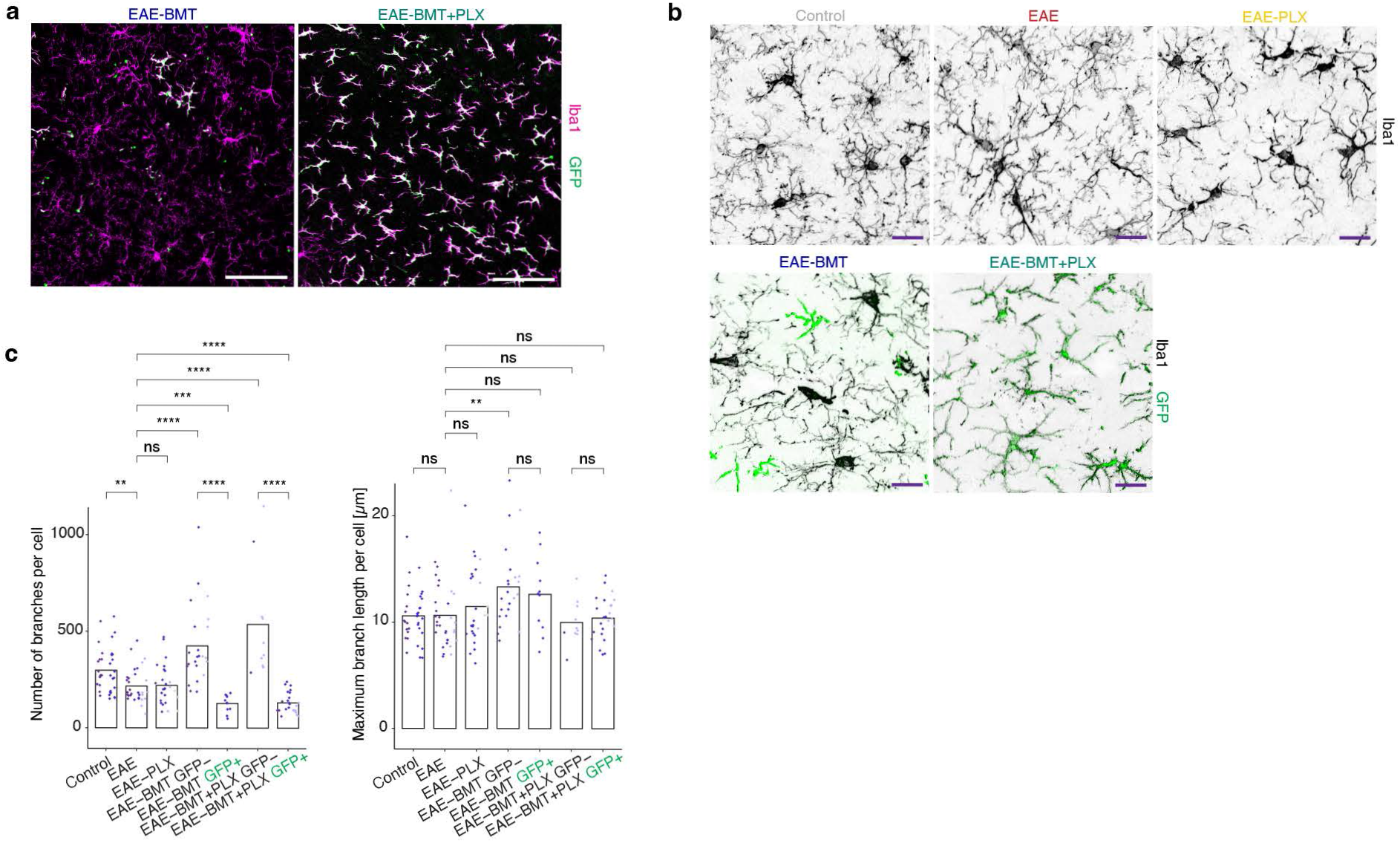
(**a**) Representative images of GFP+ chimerism in brain myeloid cells. (**b**) Representative immunofluorescent images of ramified myeloid cells in the brain. Brightness/contrast was adjusted individually per image for morphological assessment despite differing Iba1 intensity between conditions. Scale bar is 20µm. (**c**) Morphological analysis of ramified myeloid cells of the cortex in different conditions. Bars represent the group mean. Each dot represents one cell, dot colors represent different animals. Mann–Whitney U test; ns: p>0.05, *: p≤0.05, **: p≤0.01, ***: p≤0.001, ****: p≤0.0001.

**Extended Data Figure 6.**
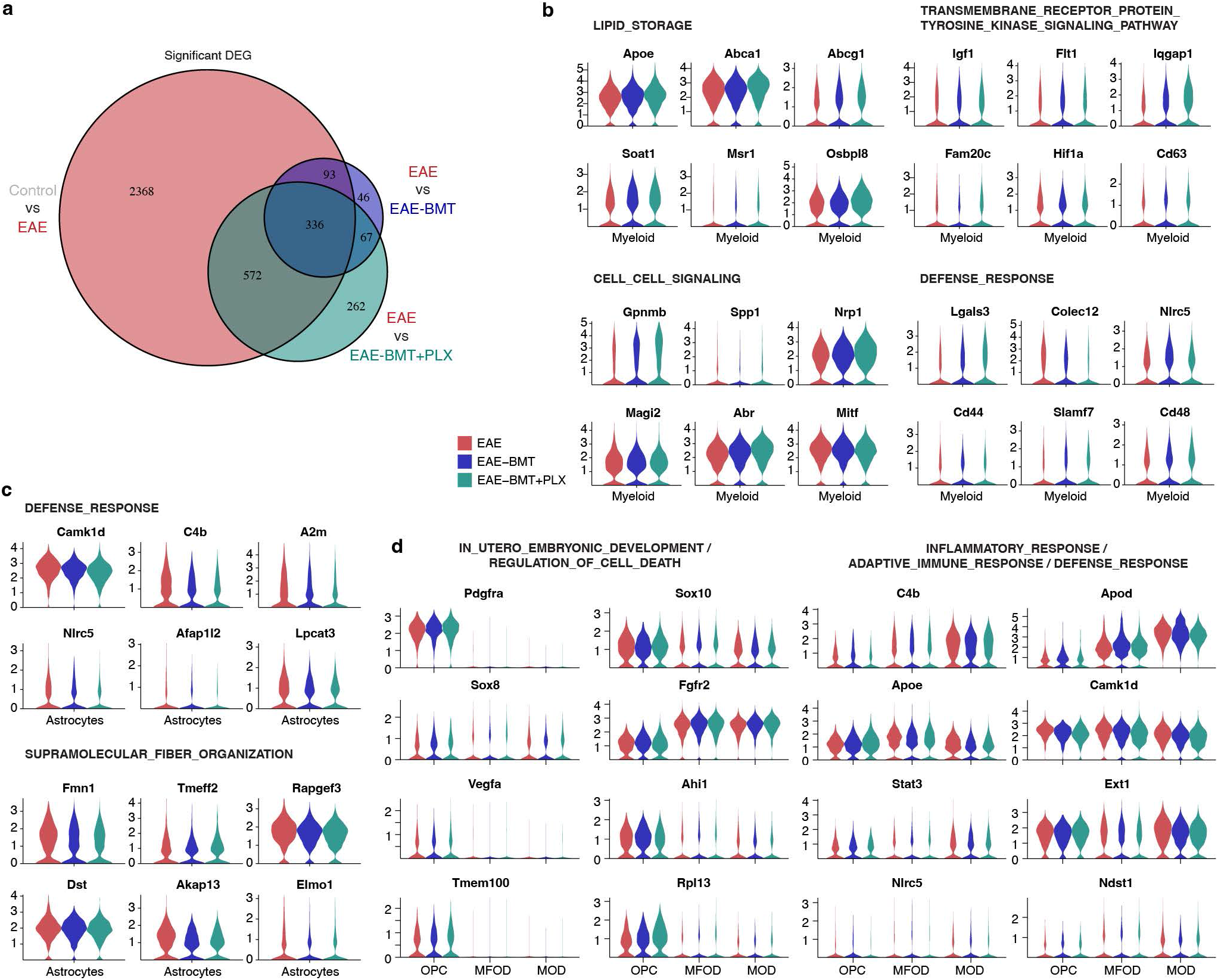
(**a**) Venn diagram showing significant DEGs (padj <0.05) of the whole NucSeq dataset between different conditions. (**b**)-(**d**) Selected genes of certain EAE-enriched pathways in the context of BMT and BMT+PLX for myeloid cells (**b**), astrocytes (**c**), and OLC (**d**).

**Extended Data Figure 7.**
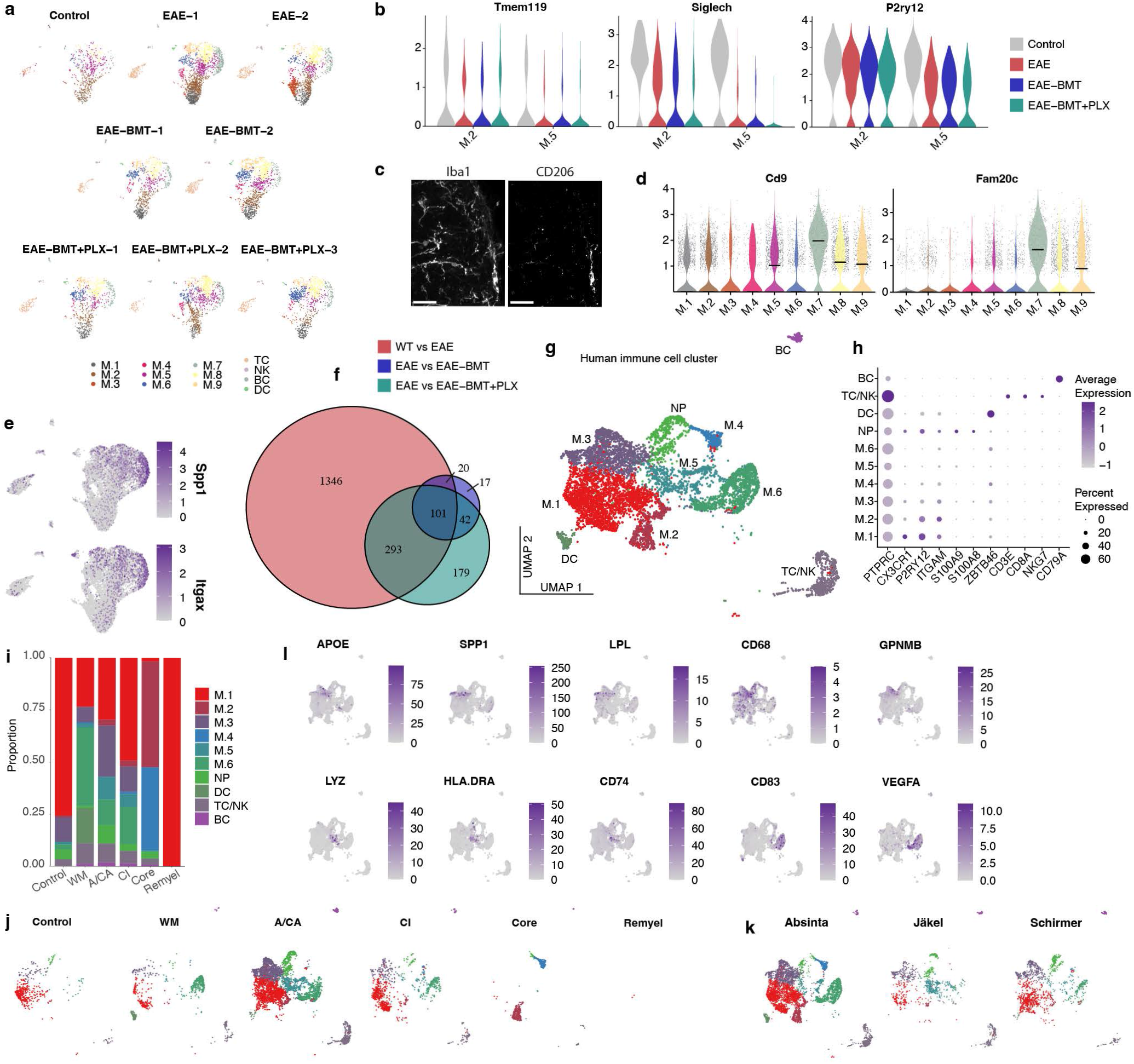
(**a**) UMAP of the immune cell cluster split by condition. (**b**) Microglia marker genes are shown for cluster M.2 and M.5 for different conditions. (**c**) Immunostaining of the white matter in a *Control* spinal cord demonstrating a CD206 positive border-associated macrophage. Scale bar is 30µm. (**d**) Expression of additional DAM (disease-associated microglia) genes associated with subcluster M.7. (**e**) Co- expression of *Spp1* and *Itgax* in cluster M.7 is demonstrated. Dots are ordered towards the front based on expression. (**f**) Venn diagram showing significant DEGs (padj <0.05) of the myeloid subcluster between different conditions. (**g**) UMAP of the immune cell cluster of the human dataset, representing 7822 cells. BC, B cells. DC, dendritic cells. M, myeloid. NP, neutrophils. TC/NK, T cells/natural killer cells. (**h**) Canonical marker genes of immune cells. (**i**) Distribution of immune cell clusters between control and MS conditions. (**j**) UMAPs split by control and MS conditions. (**k**) UMAPs split by original dataset. (**l**) Marker genes of clusters M.3 (top row), M.5 (bottom row, left), and M.6 (bottom row, right).

**Extended Data Figure 8.**
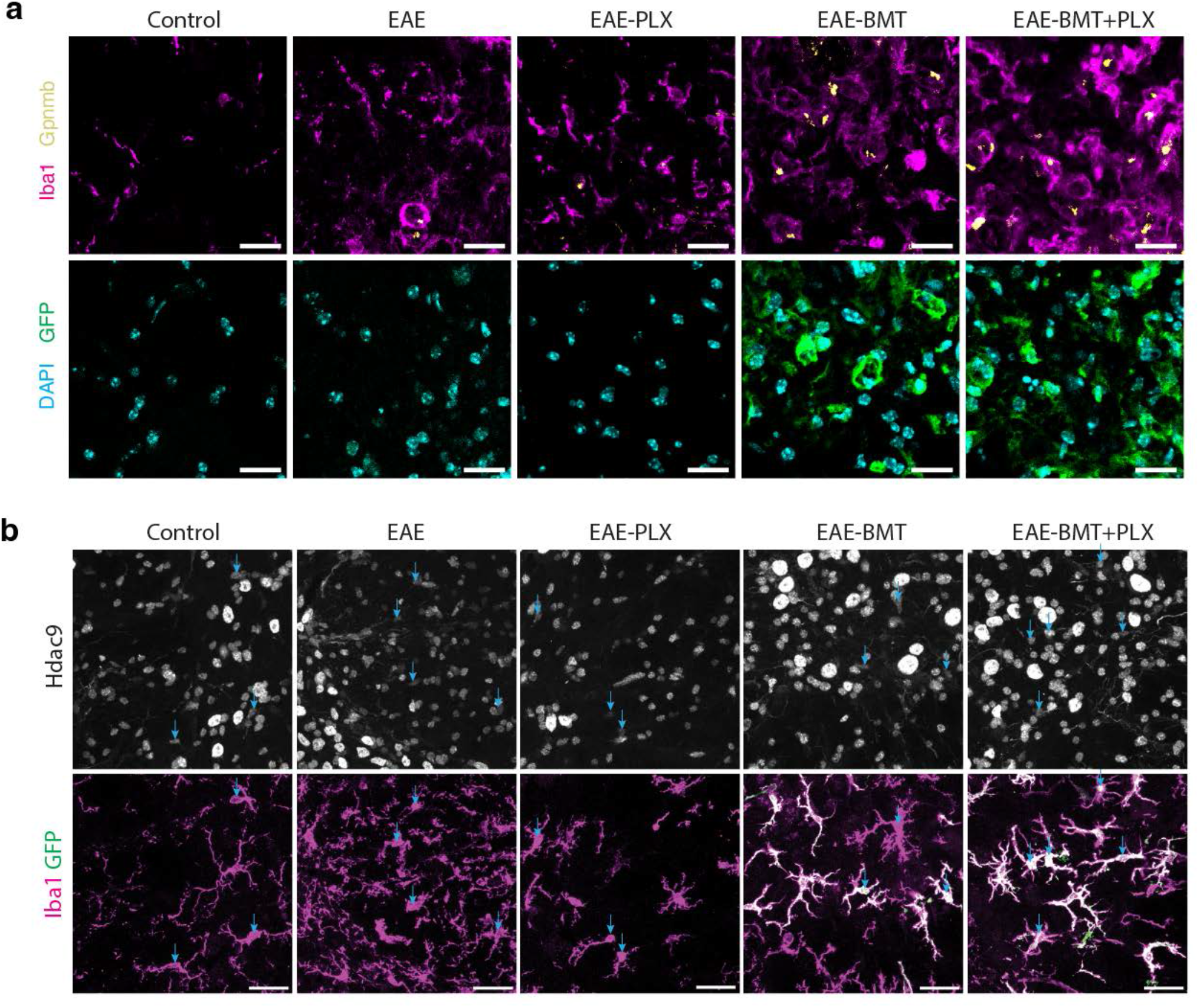
(**a**) Immunofluorescent stain for Gpnmb in the spinal cord white matter. Scale bar is 20µm. (**b**) Immunofluorescence images of the grey/white matter border in the spinal cord. Neuronal nuclei show bright Histone deacetylase 9 (Hdac9) signal. Light blue arrows indicate colocalization of myeloid cells and Hdac9 positive nuclei. Scale bar is 30µm.

## References

1. Wallin, M. T. et al. Global, regional, and national burden of multiple sclerosis 1990–2016: a systematic analysis for the Global Burden of Disease Study 2016. The Lancet Neurology 18, 269–285 (2019).

2. Miller, A. E. et al. Autologous Hematopoietic Stem Cell Transplant in Multiple Sclerosis: Recommendations of the National Multiple Sclerosis Society. JAMA Neurol 78, 241 (2021).

3. Noseworthy, J. H., Lucchinetti, C., Rodriguez, M. & Weinshenker, B. G. Multiple Sclerosis. N Engl J Med 343, 938–952 (2000).

4. Yong, H. Y. F. & Yong, V. W. Mechanism-based criteria to improve therapeutic outcomes in progressive multiple sclerosis. Nat Rev Neurol 18, 40–55 (2022).

5. Muraro, P. A. et al. Autologous haematopoietic stem cell transplantation for treatment of multiple sclerosis. Nat Rev Neurol 13, 391–405 (2017).

6. Mancardi, G. L. et al. Autologous hematopoietic stem cell transplantation in multiple sclerosis. 10 (2015).

7. Nash, R. A. et al. High-Dose Immunosuppressive Therapy and Autologous Hematopoietic Cell Transplantation for Relapsing-Remitting Multiple Sclerosis (HALT-MS): A 3-Year Interim Report. JAMA Neurol 72, 159 (2015).

8. Atkins, H. L. et al. Immunoablation and autologous haemopoietic stem-cell transplantation for aggressive multiple sclerosis: a multicentre single-group phase 2 trial. The Lancet 388, 576–585 (2016).

9. Moore, J. J. et al. Prospective phase II clinical trial of autologous haematopoietic stem cell transplant for treatment refractory multiple sclerosis. J Neurol Neurosurg Psychiatry 90, 514–521 (2019).

10. Burt, R. K. et al. Effect of Nonmyeloablative Hematopoietic Stem Cell Transplantation vs Continued Disease-Modifying Therapy on Disease Progression in Patients With Relapsing- Remitting Multiple Sclerosis: A Randomized Clinical Trial. JAMA 321, 165 (2019).

11. Muraro, P. A. et al. Thymic output generates a new and diverse TCR repertoire after autologous stem cell transplantation in multiple sclerosis patients. Journal of Experimental Medicine 201, 805–816 (2005).

12. Muraro, P. A. et al. T cell repertoire following autologous stem cell transplantation for multiple sclerosis. J. Clin. Invest. 124, 1168–1172 (2014).

13. Massey, J. et al. Haematopoietic Stem Cell Transplantation Results in Extensive Remodelling of the Clonal T Cell Repertoire in Multiple Sclerosis. Front. Immunol. 13, 798300 (2022).

14. Ruder, J. et al. Dynamics of T cell repertoire renewal following autologous hematopoietic stem cell transplantation in multiple sclerosis. SCIENCE TRANSLATIONAL MEDICINE 17 (2022).

15. Sailor, K. A. et al. Hematopoietic stem cell transplantation chemotherapy causes microglia senescence and peripheral macrophage engraftment in the brain. Nat Med 28, 517–527 (2022).

16. Shibuya, Y. et al. Treatment of a genetic brain disease by CNS-wide microglia replacement. Sci Transl Med 14, eabl9945 (2022).

17. Gold, R. Understanding pathogenesis and therapy of multiple sclerosis via animal models: 70 years of merits and culprits in experimental autoimmune encephalomyelitis research. Brain 129, 1953–1971 (2006).

18. Psenicka, M. W., Smith, B. C., Tinkey, R. A. & Williams, J. L. Connecting Neuroinflammation and Neurodegeneration in Multiple Sclerosis: Are Oligodendrocyte Precursor Cells a Nexus of Disease? Front. Cell. Neurosci. 15, 654284 (2021).

19. Williams, J. L. et al. Astrocyte-T cell crosstalk regulates region-specific neuroinflammation. Glia 68, 1361–1374 (2020).

20. Meijer, M. et al. Epigenomic priming of immune genes implicates oligodendroglia in multiple sclerosis susceptibility. Neuron 110, 1193–1210.e13 (2022).

21. Ajami, B. et al. Single-cell mass cytometry reveals distinct populations of brain myeloid cells in mouse neuroinflammation and neurodegeneration models. Nat Neurosci 21, 541–551 (2018).

22. Guerrero, B. L. & Sicotte, N. L. Microglia in Multiple Sclerosis: Friend or Foe? Frontiers in Immunology 11, (2020).

23. Miron, V. E. et al. M2 microglia and macrophages drive oligodendrocyte differentiation during CNS remyelination. Nat Neurosci 16, 1211–1218 (2013).

24. Lampron, A. et al. Inefficient clearance of myelin debris by microglia impairs remyelinating processes. Journal of Experimental Medicine 212, 481–495 (2015).

25. Falcão, A. M. et al. Disease-specific oligodendrocyte lineage cells arise in multiple sclerosis. Nat Med 24, 1837–1844 (2018).

26. Jordão, M. J. C. et al. Single-cell profiling identifies myeloid cell subsets with distinct fates during neuroinflammation. Science 363, eaat7554 (2019).

27. DeTomaso, D. et al. Functional interpretation of single cell similarity maps. Nat Commun 10, 4376 (2019).

28. Hahn, O. et al. A spatiotemporal map of the aging mouse brain reveals white matter tracts as vulnerable foci. http://biorxiv.org/lookup/doi/10.1101/2022.09.18.508419 (2022) doi:10.1101/2022.09.18.508419.

29. International Multiple Sclerosis Genetics Consortium et al. Multiple sclerosis genomic map implicates peripheral immune cells and microglia in susceptibility. Science 365, eaav7188 (2019).

30. Absinta, M. et al. A lymphocyte–microglia–astrocyte axis in chronic active multiple sclerosis. Nature 597, 709–714 (2021).

31. Schirmer, L. et al. Neuronal vulnerability and multilineage diversity in multiple sclerosis. Nature 573, 75–82 (2019).

32. Jäkel, S. et al. Altered human oligodendrocyte heterogeneity in multiple sclerosis. Nature 566, 543–547 (2019).

33. Hohsfield, L. A. et al. Effects of long-term and brain-wide colonization of peripheral bone marrow-derived myeloid cells in the CNS. Journal of Neuroinflammation 17, 279 (2020).

34. Xu, Z. et al. Efficient Strategies for Microglia Replacement in the Central Nervous System. Cell Reports 32, 108041 (2020).

35. Cronk, J. C. et al. Peripherally derived macrophages can engraft the brain independent of irradiation and maintain an identity distinct from microglia. Journal of Experimental Medicine 215, 1627–1647 (2018).

36. Van Hove, H. et al. A single-cell atlas of mouse brain macrophages reveals unique transcriptional identities shaped by ontogeny and tissue environment. Nat Neurosci 22, 1021– 1035 (2019).

37. Bennett, F. C. et al. A Combination of Ontogeny and CNS Environment Establishes Microglial Identity. Neuron 98, 1170–1183.e8 (2018).

38. Hammond, T. R. et al. Single-Cell RNA Sequencing of Microglia throughout the Mouse Lifespan and in the Injured Brain Reveals Complex Cell-State Changes. Immunity 50, 253–271.e6 (2019).

39. Barrat, F. J., Crow, M. K. & Ivashkiv, L. B. Interferon target-gene expression and epigenomic signatures in health and disease. Nat Immunol 20, 1574–1583 (2019).

40. Keren-Shaul, H. et al. A Unique Microglia Type Associated with Restricting Development of Alzheimer’s Disease. Cell 169, 1276–1290.e17 (2017).

41. Jaitin, D. A. et al. Lipid-Associated Macrophages Control Metabolic Homeostasis in a Trem2- Dependent Manner. Cell 178, 686–698.e14 (2019).

42. Shen, X., Qiu, Y., Wight, A. E., Kim, H.-J. & Cantor, H. Definition of a mouse microglial subset that regulates neuronal development and proinflammatory responses in the brain. Proceedings of the National Academy of Sciences 119, e2116241119 (2022).

43. Schroeter, M., Zickler, P., Denhardt, D. T., Hartung, H.-P. & Jander, S. Increased thalamic neurodegeneration following ischaemic cortical stroke in osteopontin-deficient mice. Brain 129, 1426–1437 (2006).

44. Tagliabracci, V. S. et al. A Single Kinase Generates the Majority of the Secreted Phosphoproteome. Cell 161, 1619–1632 (2015).

45. Masuda, T. et al. Spatial and temporal heterogeneity of mouse and human microglia at single-cell resolution. Nature 566, 388–392 (2019).

46. Abel, A. M. et al. IQGAP1: Insights into the function of a molecular puppeteer. Molecular Immunology 65, 336–349 (2015).

47. Hüttenrauch, M. et al. Glycoprotein NMB: a novel Alzheimer’s disease associated marker expressed in a subset of activated microglia. Acta Neuropathologica Communications 6, 108 (2018).

48. Liguori, M. et al. The soluble glycoprotein NMB (GPNMB) produced by macrophages induces cancer stemness and metastasis via CD44 and IL-33. Cell Mol Immunol 18, 711–722 (2021).

49. Haimon, Z. et al. Cognate microglia–T cell interactions shape the functional regulatory T cell pool in experimental autoimmune encephalomyelitis pathology. Nat Immunol (2022) doi:10.1038/s41590-022-01360-6.

50. Casamassa, A. et al. Ncx3 gene ablation impairs oligodendrocyte precursor response and increases susceptibility to experimental autoimmune encephalomyelitis. Glia 64, 1124–1137 (2016).

51. Lee, H. K. et al. Daam2-PIP5K Is a Regulatory Pathway for Wnt Signaling and Therapeutic Target for Remyelination in the CNS. Neuron 85, 1227–1243 (2015).

52. Escartin, C. et al. Reactive astrocyte nomenclature, definitions, and future directions. Nat Neurosci 24, 312–325 (2021).

53. Huang, Y. et al. Repopulated microglia are solely derived from the proliferation of residual microglia after acute depletion. Nat Neurosci 21, 530–540 (2018).

54. Elmore, M. R. P. et al. Colony-Stimulating Factor 1 Receptor Signaling Is Necessary for Microglia Viability, Unmasking a Microglia Progenitor Cell in the Adult Brain. Neuron 82, 380–397 (2014).

55. Hwang, D. et al. CSF-1 maintains pathogenic but not homeostatic myeloid cells in the central nervous system during autoimmune neuroinflammation. Proceedings of the National Academy of Sciences 119, e2111804119 (2022).

56. Hagan, N. et al. CSF1R signaling is a regulator of pathogenesis in progressive MS. Cell Death Dis 11, 1–25 (2020).

57. Nissen, J. C., Thompson, K. K., West, B. L. & Tsirka, S. E. Csf1R inhibition attenuates experimental autoimmune encephalomyelitis and promotes recovery. Experimental Neurology 307, 24–36 (2018).

58. Wlodarczyk, A. et al. CSF1R Stimulation Promotes Increased Neuroprotection by CD11c+ Microglia in EAE. Front. Cell. Neurosci. 12, 523 (2019).

59. Montilla, A. et al. Microglia and meningeal macrophages depletion delay the onset of experimental autoimmune encephalomyelitis. 2022.06.10.495612 Preprint at https://doi.org/10.1101/2022.06.10.495612 (2022).

60. Tanabe, S., Saitoh, S., Miyajima, H., Itokazu, T. & Yamashita, T. Microglia suppress the secondary progression of autoimmune encephalomyelitis. Glia 67, 1694–1704 (2019).

61. Lamb, Y. N. Pexidartinib: First Approval. Drugs 79, 1805–1812 (2019).

62. Red Blood Cell Lysis Buffer. Cold Spring Harb Protoc 2006, pdb.rec390 (2006).

63. Hahn, O. et al. CoolMPS for robust sequencing of single-nuclear RNAs captured by droplet- based method. Nucleic Acids Research 49, e11–e11 (2021).

64. Liu, S., Lockhart, J. R., Fontenard, S., Berlett, M. & Ryan, T. M. Mapping the Chromosomal Insertion Site of the GFP Transgene of UBC-GFP Mice to the MHC Locus. J.I. 204, 1982–1987 (2020).

65. Stuart, T. et al. Comprehensive Integration of Single-Cell Data. Cell 177, 1888–1902.e21 (2019).

66. Hafemeister, C. & Satija, R. Normalization and variance stabilization of single-cell RNA-seq data using regularized negative binomial regression. Genome Biology 20, 296 (2019).

67. Russ, D. E. et al. A harmonized atlas of mouse spinal cord cell types and their spatial organization. Nat Commun 12, 5722 (2021).

68. Blum, J. A. et al. Single-cell transcriptomic analysis of the adult mouse spinal cord reveals molecular diversity of autonomic and skeletal motor neurons. Nat Neurosci 24, 572–583 (2021).

69. Zeisel, A. et al. Molecular Architecture of the Mouse Nervous System. Cell 174, 999–1014.e22 (2018).

70. Marques, S. et al. Oligodendrocyte heterogeneity in the mouse juvenile and adult central nervous system. Science 352, 1326–1329 (2016).

71. Finak, G. et al. MAST: a flexible statistical framework for assessing transcriptional changes and characterizing heterogeneity in single-cell RNA sequencing data. Genome Biology 16, 278 (2015).

72. Korotkevich, G. et al. Fast gene set enrichment analysis. 060012 Preprint at https://doi.org/10.1101/060012 (2021).

73. Speir, M. L. et al. UCSC Cell Browser: visualize your single-cell data. Bioinformatics 37, 4578–4580 (2021).

